# Genomically recoded *Escherichia coli* with optimized functional phenotypes

**DOI:** 10.1101/2024.08.29.610322

**Authors:** Colin Hemez, Kyle Mohler, Felix Radford, Jack Moen, Jesse Rinehart, Farren J. Isaacs

## Abstract

Genomically recoded organisms hold promise for many biotechnological applications, but they may exhibit substantial fitness defects relative to their non-recoded counterparts. We used targeted metabolic screens, genetic analysis, and proteomics to identify the origins of fitness impairment in a model recoded organism, *Escherichia coli* C321.ΔA. We found that defects in isoleucine biosynthesis and release factor activity, caused by mutations extant in all K-12 lineage strains, elicited profound fitness impairments in C321.ΔA, suggesting that genome recoding exacerbates suboptimal traits present in precursor strains. By correcting these and other C321.ΔA-specific mutations, we engineered C321.ΔA strains with doubling time reductions of 17% and 42% in rich and minimal medium, respectively, compared to ancestral C321. Strains with improved growth kinetics also demonstrated enhanced ribosomal non-standard amino acid incorporation capabilities. Proteomic analysis indicated that C321.ΔA lacks the ability to regulate essential amino acid and nucleotide biosynthesis pathways, and that targeted mutation reversion restored regulatory capabilities. Our work outlines a strategy for the rapid and precise phenotypic optimization of genomically recoded organisms and other engineered microbes.

## Introduction

Genome recoding—the replacement of all occurrences of a codon within an organism’s genome with a synonymous codon—constitutes a promising strategy for constructing strains with broad utility in biomedicine, environmental engineering, and industrial biosynthesis^1–3^. One model recoded organism is C321.ΔA (hereafter abbreviated C321), a K-12 lineage *Escherichia coli* strain that lacks all 321 instances of UAG stop codons as well as release factor 1 (RF1), which catalyzes the ribosomal release of peptides encoded by UAG- and UAA-ending mRNAs^4^. C321 shows enhanced ribosomal nonstandard amino acid (nsAA) incorporation^5^, broad resistance to horizontal gene transfer^6,7^, and robust biocontainment when engineered to depend on exogenously-supplied nsAAs^8,9^. However, C321 and other genomically recoded strains^10,11^ show substantial fitness defects, as measured by growth rate in both rich and minimal media, compared to their non-recoded counterparts. This limits their utility in numerous research and industrial settings. For instance, existing recoded strains could exhibit impairment for synthesizing biochemicals and biologic medications at industrial scales, wherein the robust growth of host strains is a highly desirable trait^12^, despite their expanded biosynthetic capabilities.

The origins of the phenotypic impairments observed in genomically recoded strains remain unclear. Whole-genome codon reassignment may carry intrinsic fitness costs by shifting the balance between codon abundance and codon-associated translational machinery (tRNAs for proteinogenic codons and release factors for stop codons)^13^ or by inducing poorly-tolerated changes in the transcriptional or translational activity of crucial genes^14^. Alternatively, secondary mutations acquired over the course of recoded strain construction could be responsible for phenotypic impairments. The nature of fitness reductions seen in existing genomically recoded strains has implications for more extensive codon reassignment efforts that are currently underway in *E. coli*^10,11^ and other organisms^15–17^. If recoding introduces intrinsic fitness costs, there may be a ceiling to the amount of codon reassignment that an organism can tolerate; if secondary mutations are the primary cause of the observed fitness defects, then codon reassignment may be a broadly generalizable strategy for engineering synthetic organisms with diverse capabilities. Elucidating the precise genetic causes of phenotypic impairment in C321 would thus inform design strategies for ongoing and upcoming codon reassignment campaigns.

Past studies have sought to reduce the growth impairments of C321 using targeted mutagenesis and adaptive evolution. Kuznetsov et al. used multiplex genome engineering (MAGE)^18,19^ to correct a subset of the 360 secondary point mutations that arose during the construction of C321^4,20^. This modified strain, C321.ΔA.opt (abbreviated C321.OPT), showed a ∼30% improvement in growth rate relative to ancestral C321 in rich medium (LB). Wannier et al. passaged C321 in glucose-supplemented minimal medium (M9) for ∼1100 generations and isolated a strain, C321.ΔA.M9adapted (abbreviated C321.M9), that showed a ∼200% improvement in growth rate relative to ancestral C321 in M9^21^. However, the methodologies used to improve phenotype in these studies have several disadvantages. Targeted mutagenesis enables the precise reversion of desired mutations, but is limited in scope because researchers must select mutations for reversion *a priori*. Adaptive evolution does enable mutational sampling at the whole-genome scale, but runs the risk of selecting for a phenotype that is optimal under evolution conditions at the expense of other desired traits due to the acquisition of off-target and compensatory mutations^22^. Neither strategy identifies the underlying genetic cause(s) of phenotype impairment.

In this study, we build on past strain optimization efforts by precisely identifying and rectifying growth-impairing mutations in C321 using metabolic profiling, genetic analysis, and proteomic analysis. We hypothesized that mutations within genes encoding essential biosynthetic pathways would have a high likelihood of contributing to growth impairments in C321. By culturing C321 in minimal medium supplemented with combinations of amino acids, nucleotides, essential cofactors, and trace minerals, we identified metabolites whose omission reduces C321 growth. We found a deficiency in isoleucine biosynthesis that has a strong deleterious impact on growth and that is caused by a frameshifting two-nucleotide deletion in the *ilvG* gene common to all K-12 *E. coli* lineage strains^23^. Reverting this deletion dramatically improved C321 strain fitness, and installing additional fitness-improving mutations to *ilvG*-reverted strains led to C321 variants with superior growth kinetics and enhanced nsAA incorporation in both rich and minimal media.

We also performed comparative proteomics on non-recoded *E. coli* strain EcNR2, C321, and our fitness-enhanced C321 derivatives to characterize the metabolic pathways of recoded strains in detail. Compared against its non-recoded counterpart, C321 exhibited numerous deficiencies in metabolic pathways involving amino acid, nucleotide, and siderophore biosynthesis. Many of these deficiencies were corrected in enhanced C321 variants, suggesting that the small number of mutations targeted for reversion have far-ranging phenotypic effects. Our findings indicate that preexisting metabolic deficiencies play strong roles in impairing the fitness of genomically recoded strains, and that fitness impairments associated with codon reassignment may be low relative to fitness impairments caused by secondary mutation acquisition. The strategy outlined in this study enables precise and rapid phenotype optimization following genome-scale codon reassignment and could be applied to other extensively engineered microbes.

## Results

### C321 is growth impaired in both rich and minimal media

We compared the growth of C321 against ancestral K-12 strains and non-K-12 lineage strains (**Supplementary table 1**). C321 shows growth impairments when compared against its *E. coli* K-12 ancestors EcNR2^19^, MG1655, and the non-K-12 strain BL21 in both minimal (**Figure 1A**) and rich (**Supplementary data figure 1**) medium. C321 shows more substantial growth impairments in minimal medium relative to rich medium: the doubling time of the strain is 70% greater than that of EcNR2 in minimal medium (126 ± 2 min vs. 75 ± 0.8 min; **Figure 1B**) and is 40% greater than that of EcNR2 in rich medium (43.5 ± 1 min vs. 30.3 ± 0.1 min; **Figure 1C**).

**Figure 1.**
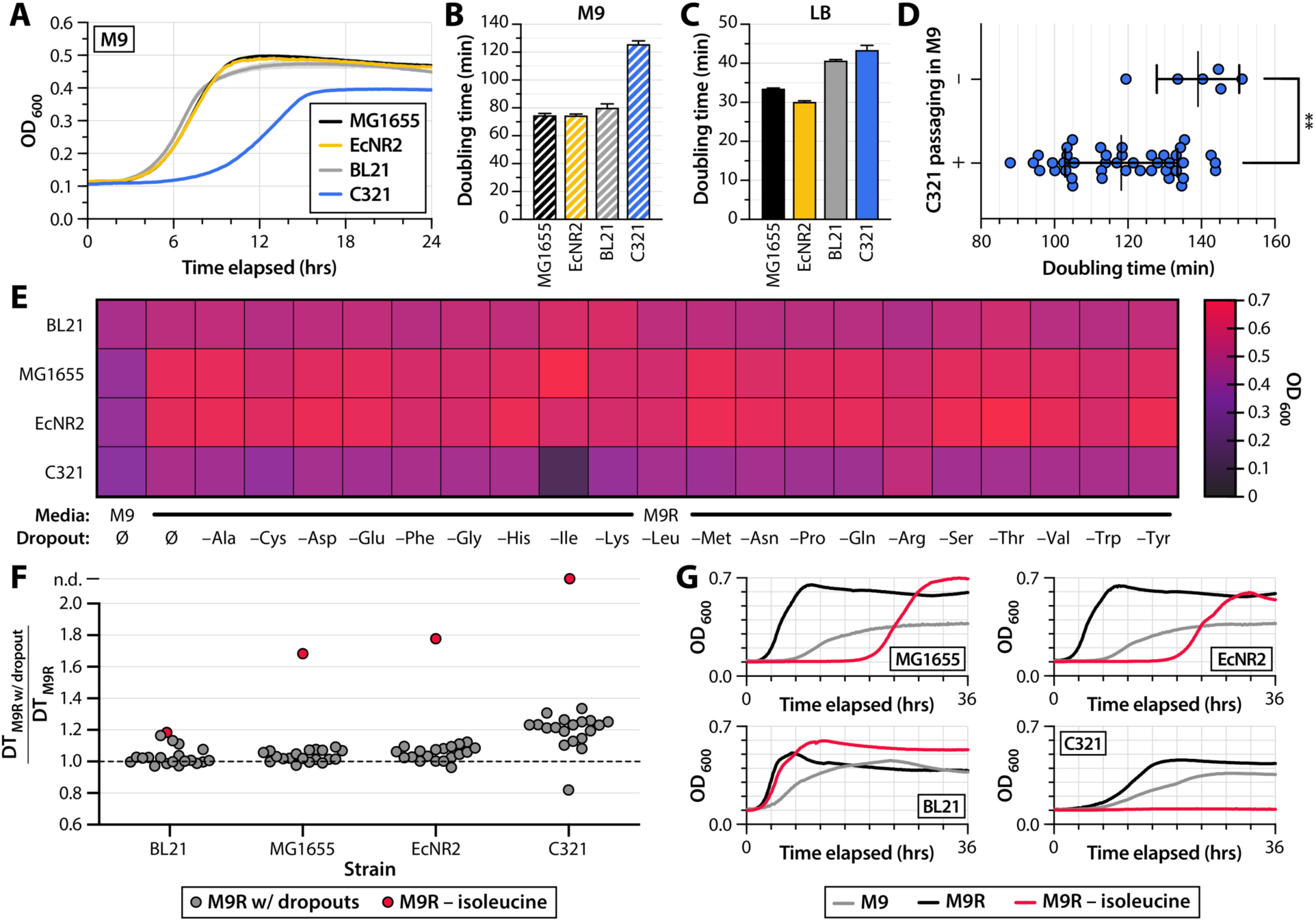
C321 shows growth impairments in rich and minimal media. (**A**) Growth curves of MG1655 (black), EcNR2 (yellow), non-K12 strain BL21 (grey), and C321 (blue) in M9 minimal medium (n = 3 for all strains). (**B-C**) Doubling times of MG1655, EcNR2, BL21, and C321 in M9 minimal medium (**B**) and LB rich medium (**C**). (**D**) Doubling times of individual C321 clones with and without serial passaging in M9 for ∼53 generations. Values are normalized to the mean doubling time for clones grown without serial passaging (comparison of means by Welch’s *t*-test; p = 0.0038). (**E**) Maximum OD_600_ reached for strains grown in M9, M9 medium supplemented with 20 amino acids (M9R), or M9R dropout medium lacking one of 20 amino acids. (**F**) Doubling times of strains in M9 dropout medium. Values are normalized to the doubling time of each strain in M9R. The normalized doubling time for each strain grown in M9R with an isoleucine dropout is highlighted in red; n.d. denotes that no growth was observed. (**G**) Growth curves for strains grown in M9 (grey), M9R (black), and M9R with an isoleucine dropout (red).

We passaged cultures of C321 in M9 and LB to determine if the strain evolves faster growth rates via adaptive evolution. After approximately 53 generations, single colonies of C321 passaged in M9 showed a 15% reduction in mean doubling time compared to un-passaged single colonies (**Figure 1D**; Welch’s *t*-test for difference in means: *p* = 0.0038; F-test for difference in variances: *p* = 0.51). Single colonies of C321 passaged in LB demonstrated a more modest (5.4%) reduction in mean doubling time after 66 generations (**Supplementary data figure 2**; Welch’s *t*-test: *p* = 0.014; F-test: *p* = 0.97). The rapidly-emerging and substantial improvements in growth rate we observed after serial passaging in M9 suggested that large fitness improvements could be engineered into C321 with relatively few genetic mutations. Rather than pursue an adaptive evolution approach that could compound mutations on a mutant genotype, we were motivated to search for mutations within C321’s genome that have strongly deleterious impacts on growth rate in M9.

### A deficiency in isoleucine biosynthesis contributes to growth impairments in C321

C321’s conspicuous growth impairment in M9 compared to LB suggests a deficiency in the ability to synthesize one or greater biomolecules that are absent in minimal medium. These include amino acids, nucleobases, and essential enzyme cofactors. We reasoned that identifying the metabolic underpinnings of C321’s growth impairment would help us to identify regions of the *E. coli* metabolome that may be dysregulated in C321. Mutations in genes within these pathways would have a high likelihood of contributing to C321’s growth impairments and would be pertinent targets for reversion.

We grew C321 and ancestral strains in M9 supplemented with combinations of the 20 proteinogenic amino acids (20AA), nucleobases (ACGU), and/or essential cofactors and trace metals (vitamin mix) (supplements are detailed in **Supplementary table 2**). We observed profound reductions in maximum OD_600_ when C321 was grown in media lacking a 20-amino acid supplement, but not when lacking a nucleobase supplement or vitamin mix (**Supplementary data figure 3**). This suggested that a deficiency in amino acid biosynthesis contributes strongly to C321’s growth impairment in minimal media. To determine which amino acid(s) were responsible, we next grew C321 and ancestral strains in M9 media supplemented with all 20 amino acids (designated M9 rich medium; M9R) and in M9R dropout media lacking one of each amino acid (**Figure 1E**; growth curves shown in **Supplementary data figure 4**). C321 did not grow in M9R lacking isoleucine.

Whereas the omission of one amino acid from M9R led to modest, if any, increases in doubling time for most dropouts, the omission of isoleucine from M9R led to increases in doubling time for all strains (**Figure 1F**). Notably, increases in doubling time for isoleucine dropout medium were higher for K-12 lineage strains MG1655 and EcNR2 (70% and 80% increases relative to growth in M9R, respectively) than they were for BL21 (20% increase relative to growth in M9R). MG1655 and EcNR2 also showed profound delays in time taken to reach mid-log phase after inoculation compared to BL21 (**Figure 1G**). These observations raised the possibility that the growth defect observed in C321 is attributable to dysregulated isoleucine biosynthesis caused by a mutation that is also present in K-12 ancestral strains. As has been noted in early published *E. coli* complete genome sequences^23^, all K-12 lineage strains, including C321, have two-nucleotide deletion—Δ(A979-T980)—in the isoleucine biosynthesis gene *ilvG*, which encodes the catalytic subunit of acetolactate synthase. The frameshift introduced by this deletion places a TGA stop codon one residue downstream of its occurrence, effectively truncating 221 C-terminal residues from the IlvG protein.

### K-12 lineage mutations targeted for reversion in C321

Prior work demonstrated that correcting the Δ(A979-T980) frameshift in *ilvG* eliminates the metabolic oscillatory behavior observed in K-12 strains and reduces growth impairments in media containing glucose as a sole carbon source^24^. We hypothesized that the same correction in C321 would alleviate the strain’s growth defects, and we sought to identify additional K-12 mutations for targeted reversion in C321. We suspected that an alanine-to-threonine substitution in position 246 of UAA-/UGA-release factor 2, encoded by *prfB* and common to all K-12 strains, could contribute to C321’s growth impairments. PrfB.A246T variants show a reduction in the release rate of UAA codons *in vitro*^25^, and prior adaptive evolution of C321 in minimal medium demonstrated spontaneous reversion of the A246T mutation^21^.

### Reversions of K-12 lineage mutations in prfB and ilvG are sufficient to alleviate growth impairments in C321

We first investigated the role that K-12 lineage mutations played in the growth impairments of C321. We used multiplex genome engineering (MAGE)^19^ to correct mutations in *prfB* and *ilvG* in both EcNR2 and C321. EcNR2 harboring these reversions (designated EcNR2.*prfB*^+^.*ilvG*^+^) demonstrated no improvements in growth phenotype (**Figure 2A**). C321 with *prfB* and *ilvG* corrections (designated C321.*prfB*^+^.*ilvG*^+^) showed a modest growth improvement in rich medium (5.5% reduction in doubling time relative to C321) (**Figure 2B**). However, the corrections led to a near-complete recovery of growth in minimal medium, reducing doubling time relative to C321 by 37%. The doubling time of C321.*prfB*^+^.*ilvG*^+^ in minimal media is 110% that of EcNR2.

**Figure 2.**
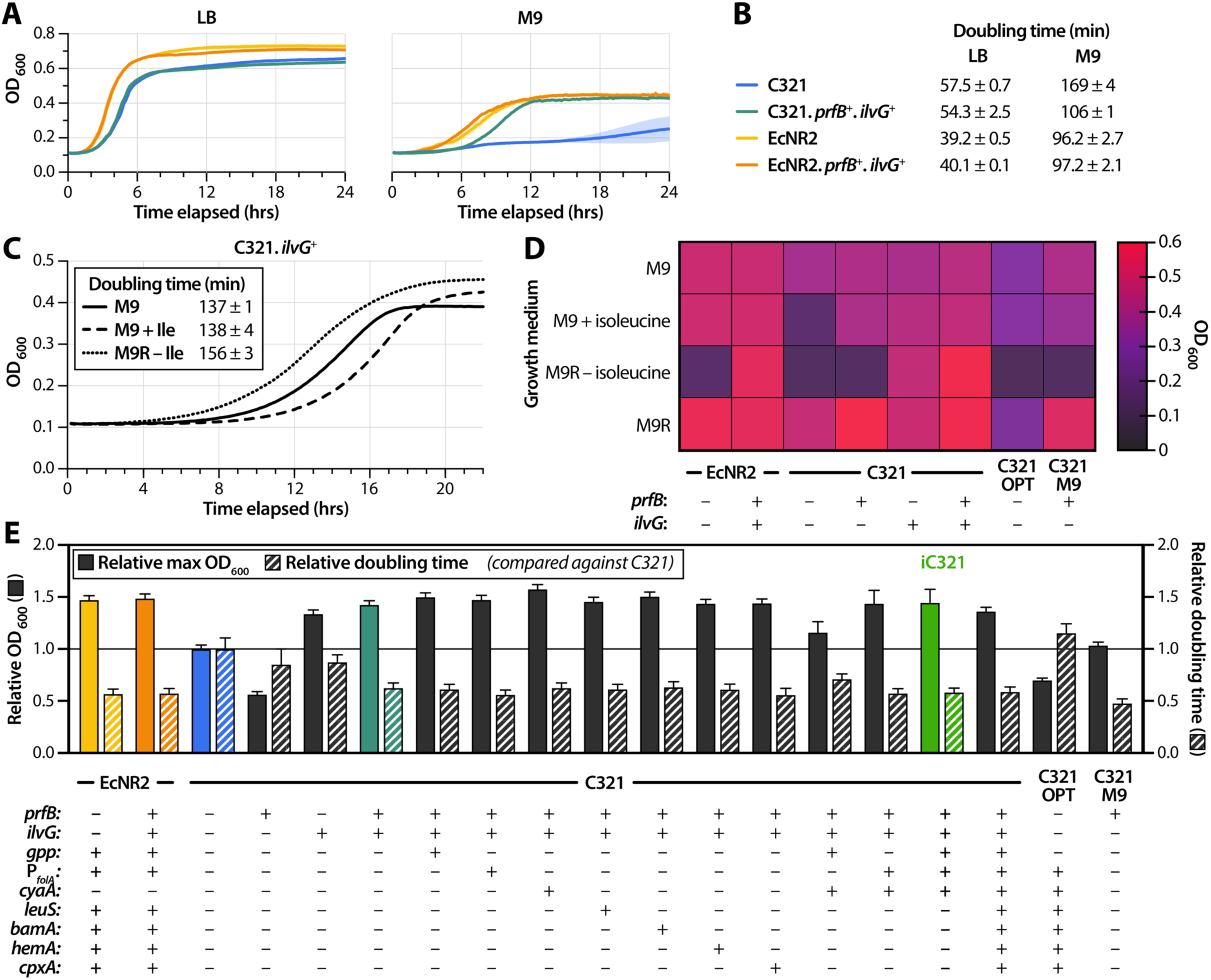
Reversion of K-12 lineage mutations alleviates growth impairments in C321. (**A**) Growth curves of EcNR2 and C321 with and without reversion of K-12 lineage mutations (*prfB* and *ilvG*) in LB (left panel) and M9 (right panel). (**B**) Doubling times of strains shown in panel A (n = 3). (**C**) Growth curves of C321.*ilvG^+^* in M9 (solid line), M9 supplemented with isoleucine (dashed line), and M9R with an isoleucine dropout (dotted line). (**D**) Maximum OD_600_ reached for strains with reversions in K-12 lineage mutations grown in M9 with a variety of amino acid supplements. C321.OPT denotes an engineered C321 variant reported by Kuznetsov et al. 2017, and C321.M9 denotes an evolved C321 variant reported by Wannier et al. 2018. (**E**) Maximum OD_600_ and doubling time relative to C321 for engineered C321 strains in M9 (*n* = 3 biological replicates). The solid line denotes the maximum OD_600_ and doubling time of C321. One-way ANOVA for comparisons between all strains is given in Supplementary table 6.

C321 with a single reversion in *ilvG* (designated C321.*ilvG*^+^) did not show an extended lag phase after inoculation when cultured in isoleucine dropout medium (**Figure 2C**, dotted line). This suggests correction of dysregulated isoleucine biosynthesis. Furthermore, C321.*ilvG*^+^ exhibited similar growth characteristics when grown in M9 without (doubling time: 137 ± 1 min) and with (doubling time: 138 ± 4 min) isoleucine supplementation (**Figure 2C**, solid and dashed lines). This indicates that restoring IlvG function is sufficient for eliminating the strain’s isoleucine dependence.

We investigated the effects of single reversions of *prfB* and *ilvG* on the growth characteristics of C321 by measuring maximum OD_600_ in minimal media and supplemented derivatives. We observed an epistatic pattern whereby *prfB* reversion only has an appreciable effect on growth if isoleucine starvation was alleviated, either through exogenous supplementation or by reversion of *ilvG* (**Figure 2D**). While *ilvG* reversion is sufficient to restore regulation of isoleucine biosynthesis, *prfB* reversion enables growth to a higher optical density in a manner unrelated to isoleucine metabolism, as C321 with a single reversion in *prfB* (designated C321.*prfB*^+^) grows more robustly than does C321 in minimal medium supplemented with all 20 amino acids (M9R). As with its recoded counterpart, EcNR2.*prfB*^+^.*ilvG*^+^ was capable of growth in M9R lacking isoleucine, indicating restoration of dysregulated isoleucine biosynthesis. We also compared the maximum OD_600_ values reached by C321.*prfB*^+^.*ilvG*^+^ against those reached by previously reported optimized C321 strains: C321.OPT and C321.M9^20,21^. C321.*prfB*^+^.*ilvG*^+^ outgrew both of these strains in all media types tested (**Figure 2D**).

### C321-specific mutations targeted for reversion in C321

While C321.*prfB*^+^.*ilvG*^+^ shows more favorable growth characteristics than other optimized C321 variants, the strain nonetheless demonstrates doubling times that are 35% slower in rich medium and 10% slower in minimal medium compared to the analogous unrecoded EcNR2.*prfB*^+^.*ilvG*^+^. We hypothesized that reversion of C321 lineage-specific mutations could further recover the remaining growth deficit in C321.*prfB*^+^.*ilvG*^+^. Analysis of single nucleotide substitutions in the protein-coding regions of the C321 genome (**Supplementary table 3**) revealed a R134H substitution in Gpp, a phosphatase that metabolizes pppGpp to ppGpp and plays a key role in mobilizing the stringent response that upregulates amino acid biosynthesis under starvation conditions^26^. Strains deficient in ppGpp show a reduced ability to regulate biosynthetic genes when grown in media lacking isoleucine^27^. We targeted six additional mutations—previously identified through targeted mutagenesis and adaptive evolution—that have been shown to recover 59% of the doubling time deficit of C321 in rich medium (**Supplementary table 4**)^20^. Targeting a limited set of seven C321-specific mutations for reversion (**Table 1**) allowed us to investigate the combinatorial effects of reversions on growth phenotype and to construct optimized strains in a minimal number of generations, thereby avoiding the accumulation of carrier mutations.

**Table 1.** Mutations targeted for correction in C321. K-12 lineage mutations are present in all K-12 derived strains, and C321 lineage mutations are unique to C321 and its derivatives.

| Site | Mutation | Lineage | Function |
| --- | --- | --- | --- |
| *ilvG* | Δ(A979-T980) | K-12 | Isoleucine biosynthesis |
| *prfB* | Ala246Thr | K-12 | UGA/UAA release factor |
| *gpp* | Arg134His | C321 | Stringent response |
| *folA* promoter | promoter -35 box | C321 | Tetrahydrofolate biosynthesis |
| *cyaA* | Pro301Leu | C321 | cAMP biosynthesis and signaling |
| *leuS* | Val613Phe | C321 | Leucine aminoacyl tRNA synthetase |
| *bamA* | Pro763Ser | C321 | Outer membrane protein assembly |
| *hemA* | Leu196Pro | C321 | Porphyrin biosynthesis |
| *cpxA* | Trp184Arg | C321 | Inner membrane stress response |

### C321-specific reversions further recover phenotype in rich and minimal media

We used MAGE to generate C321.*prfB*^+^.*ilvG*^+^ derivatives with single reversions, intermediate combinations of reversions, and all reversions of C321-specific mutations. We assessed strain fitness using 96-well microplate assays to measure maximum OD_600_ and doubling time in LB and M9, and benchmarked these metrics against those obtained for C321 (**Supplementary table 5**). All engineered variants showed modest phenotype improvements compared to the improvements demonstrated by C321.*prfB*^+^.*ilvG*^+^ in both minimal medium (**Figure 2E**) and rich medium (**Supplementary data figure 5**). One-way ANOVA for all strain-by-strain comparisons is detailed in **Supplementary table 6**.

Our optimized strains demonstrate differing degrees of phenotypic improvement compared to C321 in rich and minimal medium. C321.*prfB*^+^.*ilvG*^+^ recovers 17% of the doubling time deficit relative to EcNR2 in rich medium and 87% of the doubling time deficit relative to EcNR2 in minimal medium. We identified an intermediate strain—C321.*prfB*^+^.*ilvG*^+^ with reversions of mutations in *gpp*, the *folA* promoter, and *cyaA*—that recovers 3.4% of the doubling time deficit relative to EcNR2 in rich medium and 97% of the doubling time deficit relative to EcNR2 in minimal medium. We designated this strain iC321, an abbreviation for “improved C321,” to reflect the substantial growth rate improvements it exhibits in minimal medium.

C321.*prfB*^+^.*ilvG*^+^ with all seven C321-specific mutation reversions likewise showed a similar improvement of doubling time in M9, but reached a lower maximum OD_600_ than did iC321. iC321 has a slower doubling time in LB than both C321.*prfB*^+^.*ilvG*^+^ and C321.*prfB*^+^.*ilvG*^+^ with all seven C321-specific mutation reversions, suggesting epistasis among the alleles that confer faster growth rates in rich medium. We did not observe the same epistatic pattern for growth rates in minimal medium. C321.*prfB*^+^.*ilvG*^+^ with all seven C321-specific mutation reversions recovers 54% of the doubling time deficit relative to EcNR2 in rich medium and 95% of the doubling time deficit relative to EcNR2 in minimal medium. Our phenotype-optimized strains do not show reductions in maximum OD_600_, which we observed in the growth profiles of previously reported optimized strains C321.OPT and C321.M9 in both M9 (**Figure 2E**) and LB (**Figure 2F**).

### Phenotype-optimized C321 strains show improved proteomic profiles

C321.*prfB*^+^.*ilvG*^+^ and iC321 exhibit profound phenotypic improvements over C321 in minimal medium. To understand the physiologic basis for these improvements, we performed proteomic analysis on C321, C321.*prfB*^+^.*ilvG*^+^, iC321, and EcNR2 grown in minimal medium. We used label-free quantification (LFQ) to determine the abundances of proteins encoded by genes targeted for reversion in iC321 (**Figure 3A; Supplementary table 7**) as well as proteins encoded by genes in the isoleucine biosynthesis (*ilv*) family (**Figure 3B)**. FolA showed a much lower abundance in EcNR2-genotype strains (EcNR2 and iC321) compared to mutated strains (C321 and C321.*prfB*^+^.*ilvG*^+^). The mutation found in C321 comprises a C-to-T transition within the -35 box of the *folA* promoter (TTGACG in C321 versus TCGACG in EcNR2 and other K-12 strains). The C321 genotype is in closer alignment with the consensus sigma σ-binding sequence TTGACA; this likely increases protein abundance by raising the rate of *folA* transcription. Reverting the mutation in iC321 reduces FolA abundance to a level comparable with that measured in EcNR2.

**Figure 3.**
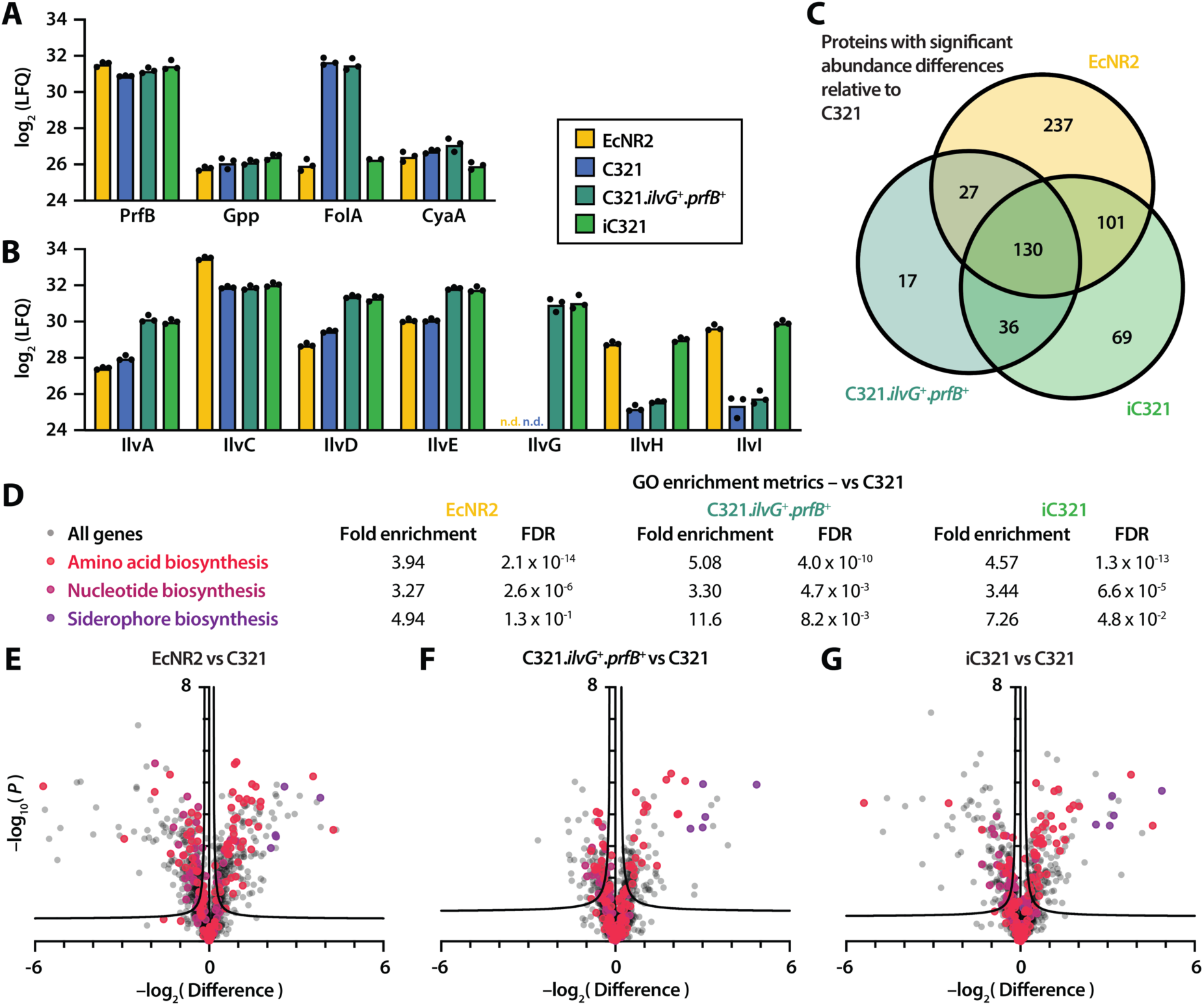
Proteomic analysis of C321 and phenotype-enhanced variants. (**A**) Label-free quantification (LFQ) intensities for peptides whose genes were targeted for mutation reversion (n = 3 biological replicates). (**B**) LFQ intensities for peptides in the *ilv* gene family; n.d. indicates that the peptide was not detected. (**C**) Set analysis of proteins with significant differences in abundance compared to C321. Regions of overlap indicate proteins with a significant difference in abundance in greater than one strain. (**D**) Gene ontology (GO) enrichment analysis for genes involved in amino acid biosynthesis (GO:0008652), nucleotide biosynthesis (GO:0009165), and siderophore biosynthesis (GO:0019290). FDR: False discovery rate. (**E-G**) Volcano plots showing whole-proteome comparisons between C321 and EcNR2 (**E**), C321.*prfB*^+^.*ilvG*^+^ (**F**), and iC321 (**G**) (mean of n = 3 biological replicates). Proteins whose genes belong to GO categories of interest are shaded according to (**D**).

Proteins encoded by *ilv* family genes showed diverse responses to mutation reversion (**Figure 3B**). Full-length IlvG was undetectable in EcNR2 and C321, but was present in both C321.*prfB*^+^.*ilvG*^+^ and iC321. Compared against C321, both C321.*prfB*^+^.*ilvG*^+^ and iC321 demonstrated increases in the abundances of IlvA/D/E. Because *ilvG* is the first gene of an operon that also encodes *ilvA*, *ilvD*, and *ilvE*, we suspect that reversion of the *ilvG* frameshift increases the expression of downstream genes through translational coupling, which has been observed within the operon between *ilvA* and *ilvD*^28^. iC321, but not C321.*prfB*^+^.*ilvG*^+^, showed increases in the abundances of IlvH/I, the functionally redundant analogues of IlvG/M. This may be attributable to the influence of intracellular (p)ppGpp concentrations on *ilvH*/*I* expression^29,30^; reversion of the C321-specific *gpp* mutation restores IlvH/I abundance to levels comparable to those measured in EcNR2. IlvC is encoded in its own single-gene operon whose activity is not known to be influenced by ppGpp signaling, and its abundance does not change appreciably in C321 strains with *ilvG* and *gpp* mutation reversions. This observation is consistent with our hypothesis that *ilvG* reversion affects the expression of downstream genes within its operon and that *gpp* reversion affects the expression of IlvI/H via modulation of the stringent response.

The genetic changes made to C321.*prfB*^+^.*ilvG*^+^ and iC321 have far-reaching effects on the strains’ proteomes. We identified all genes in EcNR2, C321.*prfB*^+^.*ilvG*^+^, and iC321 whose protein products showed significant differences in abundance compared to C321 (**Figure 3C; Supplementary table 8**). Of the 210 proteins found to have significant abundance differences in C321.*prfB*^+^.*ilvG*^+^ compared to C321, 157 (75%) were also found to be significant in EcNR2. Likewise, 231 of 336 proteins (69%) with significant abundance differences in iC321 were found to be significant in EcNR2. This suggests that the proteomic profiles of C321.*prfB*^+^.*ilvG*^+^ and iC321 are more similar to that of EcNR2 than they are to that of C321.

C321.*prfB*^+^.*ilvG*^+^ and iC321 have markedly improved biosynthetic capabilities over C321 when cultured in minimal media. We performed a gene ontology (GO) enrichment analysis^31^ on the genes with significant abundance differences compared to C321 for each strain (**Figure 3D; Supplementary tables 9-10**). We found that genes involved in amino acid biosynthesis (GO:0008652) and nucleotide biosynthesis (GO:0009165) were overrepresented in C321.*prfB*^+^.*ilvG*^+^ and iC321 as well as in EcNR2. We also found that the protein products of genes associated with siderophore biosynthesis (GO:0019290) were very strongly expressed in all three strains compared to C321 (**Figures 3E-G**, deep purple). These findings suggest that the predominant basis for C321’s growth impairment in minimal media is the dysregulation of key biosynthetic pathways as well as a deficiency in iron scavenging ability. The mutations implemented in iC321 partially restore the expression of these biosynthetic pathways, enabling higher growth rates in minimal medium.

### Enhanced nsAA incorporation with iC321

A key application for genomically recoded organisms is the efficient multisite incorporation of nonstandard amino acids (nsAAs) into proteins and protein polymers^2,4,5,32^. C321 strains lack in-frame UAG codons as well as release factor 1, which is responsible for translation termination at UAG codons. This permits the incorporation of nsAAs using aminoacyl tRNA synthetases that charge nsAAs onto tRNAs with cognate UAG codons. C321 exhibits higher specificity for multi-site nsAA incorporation compared to EcNR2^5^ in rich medium. However, C321’s fitness impairments lead to reduced total protein yields, especially in minimal medium^21^. We hypothesized that the favorable growth and proteomic characteristics of iC321 would enable high-specificity and high-yield nsAA incorporation in both rich and minimal medium.

The efficient production of nsAA-containing proteins requires a host strain with high protein production capacity and high specificity for nsAA incorporation. To assess protein production capacity, we measured non-OD-normalized fluorescence generated from the expression of ELP-GFP cassettes containing no in-frame UAG codons (**Figure 4A-B**). We compared iC321 against ancestral C321, the previously reported optimized strain C321.M9^21^, and non-recoded EcNR2 in both rich and minimal medium (we were unable to transform plasmids into C321.OPT). iC321 exhibited higher fluorescence than EcNR2 and C321.M9— but lower fluorescence than C321—in rich medium. Such differences could be due to broadly dysregulated protein synthesis pathways in C321 that are partly restored in iC321. Both iC321 and C321.M9 exhibited much higher fluorescence than C321 in minimal medium, suggesting that the mutations implemented have stronger effects on protein production capacity in minimal medium than they do in rich medium.

**Figure 4.**
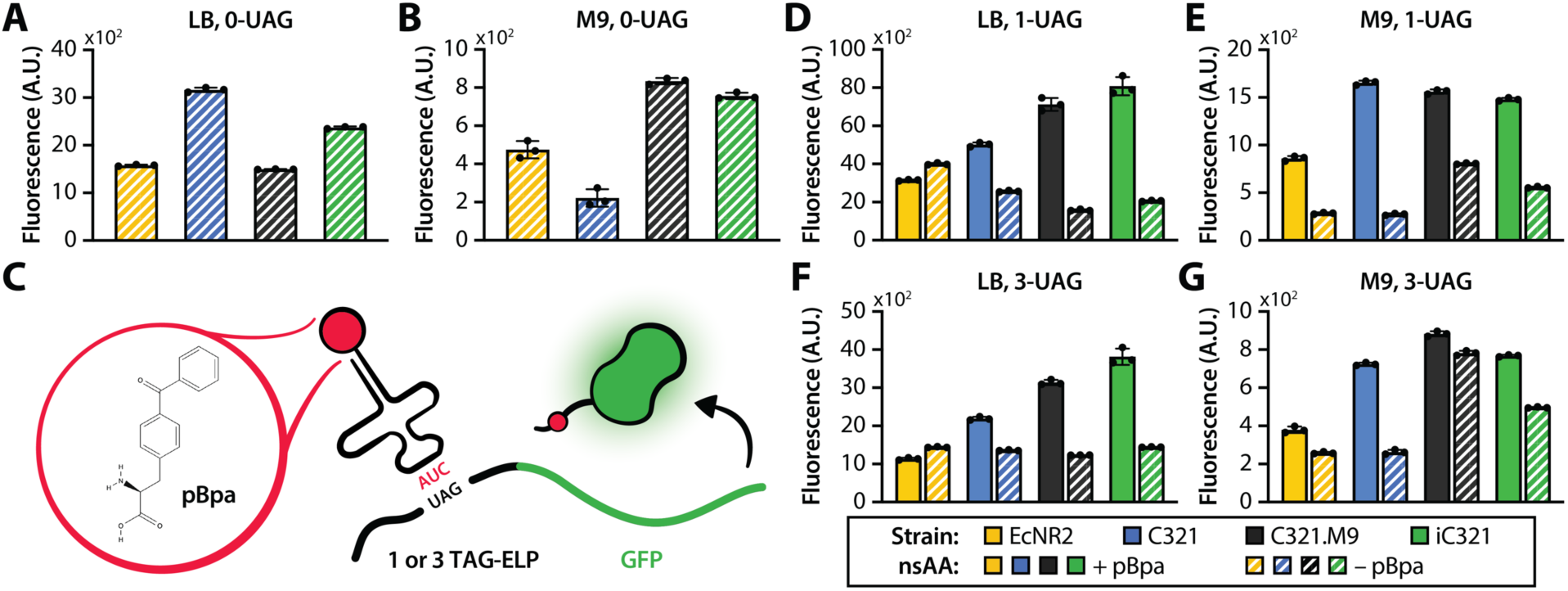
Protein production and non-standard amino acid incorporation in iC321. (**A-B**) non-OD-normalized GFP fluorescence for EcNR2, C321, and optimized C321 variants cultured in LB medium (**A**) or M9 medium (**B**) (n = 3 biological replicates). (**C**) nsAA incorporation into elastin-like polypeptide (ELP) chains fused to GFP. An orthogonal tRNA/synthetase pair encodes *para*-benzoyl-L-phenylalanine (pBpa) at UAG codons placed within the ELP sequence. (**D-F**) OD-normalized GFP fluorescence for EcNR2, C321, and optimized C321 variants expressing ELP-GFP constructs that contain 1 (**D-E**) or 3 (**E-F**) TAG codons within their coding sequences. Strains were cultured in LB medium (**D, F**) or M9 medium (**E, G**). Solid fills indicate the inclusion of 1 mM pBpa in the growth medium; striped fills indicate the omission of pBpa (*n* = 3 biological replicates).

To assess nsAA incorporation specificity, we constructed anhydrotetracycline (aTc)-inducible expression cassettes consisting of an elastin-like polypeptide (ELP) fused to the N-terminus of GFP. UAG codons placed within structurally permissive sites of the ELP chain encode nsAAs; the abundance of GFP detected via fluorescence serves as a measure of overall nsAA incorporation^5^. We expressed ELP-GFP cassettes containing one or three in-frame UAG codons alongside with a synthetase/tRNA pair that charges the photo-crosslinkable nsAA *para*-benzoyl-L-phenylalanine (pBpa) onto tRNAs with a cognate UAG codon (**Figure 4C**)^33^ and measured OD-normalized fluorescence in both rich and minimal medium (**Figures 4D-G**). We found that the fluorescence output of iC321 was higher than all other strains when expressing 1- and 3-UAG ELP-GFP in rich medium (**Figures 4D, F)**. iC321 showed nsAA incorporation activity comparable to those of C321 and C321.M9 in minimal medium (**Figures 4E, G**). Notably, fluorescence generated by the optimized strains iC321 and C321.M9 was elevated when expressing 1- and 3-UAG ELP-GFP in minimal medium without pBpa supplementation (**Figures 4E, G**, dashed bars). This suggests that the nsAA incorporation specificity of optimized strains is reduced in minimal medium, possibly due to increased rates of UAG near-cognate suppression. As has been shown previously, near-cognate suppression can trigger ribosomal rescue mechanisms that tag nascent proteins for degradation, reducing overall yields^6^.

## Discussion

Genomically recoded organisms exhibit numerous unique traits—including enhanced ribosomal nsAA incorporation, resistance to horizontal gene transfer, and biocontainment through nsAA auxotrophy—that make them appealing for a wide range of biotechnological applications^1–3^. However, C321 and other genomically recoded strains show impaired fitness compared to their non-recoded counterparts, which limits their utility. This observation has prompted unresolved questions concerning the degree to which genomic recoding imposes intrinsic fitness burdens on organisms, and thus the extent to which organisms can tolerate whole-genome codon reassignment.

To address this, we leveraged targeted metabolic screens to identify and rectify deleterious mutations in C321. We found that mutations common to all K-12 lineage strains— a two-nucleotide deletion in *ilvG* and a nonsynonymous substitution in *prfB*—play strong roles in C321’s fitness impairment in both rich and minimal media. IlvG-deficient *E. coli* strains oscillate between states of isoleucine starvation and production in media containing glucose as a sole carbon source^24^. This may be due to unchecked activity of IlvM, which functions as a regulatory subunit of full-length IlvG but can also interact with the redundant acetolactate synthase IlvI^34^. K-12 lineage *prfB* mutants show reduced UAA codon release rates *in vitro*^25^, and adaptive laboratory evolution of C321.ΔA led to the spontaneous reversion of *prfB* to a variant with higher catalytic activity^21^.

We observed an epistatic relationship for the *prfB* and *ilvG* mutations in minimal medium, whereby *prfB* correction only improved the growth phenotype of C321 in an *ilvG*-corrected background, but *ilvG* correction did improve the strain’s growth phenotype even in the absence of *prfB* correction (**Figure 2D**). We are unaware of studies that report a direct mechanistic link between PrfB and IlvG. We hypothesize that this epistatic pattern may be due to the relative demands for isoleucine and PrfB during translation—whereas virtually all proteins necessitate numerous isoleucine-charged tRNAs within their coding domains for complete synthesis, PrfB only plays a role in the termination of fully-synthesized proteins. Deficiencies in isoleucine production due to *ilvG* mutation thus mask deficiencies in translational termination due to prfB mutation. We constructed a phenotype-optimized strain, termed iC321, by correcting *ilvG* and *prfB* mutations alongside three C321-specific mutations (in *gpp*, *cyaA*, and the *folA* promoter). iC321 shows more favorable growth kinetics than previously reported optimized C321 derivatives in a range of media^20,21^, as well as an improved proteomic profile and enhanced nsAA incorporation in rich medium.

Insertion-deletion (indel) mutations are approximately 10 times less frequent per genome replication event than are single nucleotide substitutions in mismatch-repair-deficient strains like EcNR2 and C321^35^. The relative rarity of spontaneous indels helps to explain why Wannier and colleagues never observed reversion of the K-12 *ilvG* frameshift after passaging C321 in minimal medium for >1000 generations^21^. They nonetheless found that C321 exhibited appreciable reductions in doubling time after propagating in minimal medium for as few as ∼50 generations, a finding reproduced here (**Figure 1C**). Such a rapid recovery of growth impairment suggests that numerous compensatory mutations exist for *ilvG* deficiency in K-12 lineages, and that the accumulation of mutationally accessible compensatory mutations is more likely than the reversion of a single defect-inducing mutation. This pattern is consistent with past observations of compensatory evolution in diverse biological systems, including mutational trajectories taken by peptides in response to protein-protein interface-altering substitutions^36^, by mitochondrial tRNAs in response to canonical base-pair disruption^37^, and by bacterial pathogens in response to the acquisition of fitness-reducing antibiotic resistance alleles^38,39^.

Mutagenesis using MAGE can generate strains with optimized phenotypes in far fewer generations, but do so at the cost of low throughput and high expense. Kuznetsov and colleagues targeted 127 of 360 known C321-specific mutations for reversion via MAGE^20^. Assuming that the generation of each reversion is independent of all others, this corresponds to a mutational space of ∼10^38^ genotypes to explore. For comparison, there exist an estimated 10^30^ living prokaryotic cells on Earth^40^. Targeted mutagenesis at the whole-genome scale is thus unlikely to generate sufficient diversity to adequately sample all—or even a small fraction of—possible genotypes.

Strain engineering via targeted mutagenesis necessarily involves passaging cell cultures, which also permits off-target carrier and compensatory mutations to arise within lineages, albeit to a lesser degree than does adaptive evolution. The accumulation of carrier mutations during strain construction likely plays a role in reducing the versatility of recoded strains, as inadvertent adaptive evolution optimizes fitness under the conditions of strain construction at the expense of fitness in other settings. By prefacing our strain engineering efforts with a study on the metabolic basis for C321’s growth impairment, we narrowed the scope of our engineering efforts to high-impact mutations and minimized the opportunity for unintended adaptive evolution to occur. We generated C321.*prfB*^+^.*ilvG*^+^ in five MAGE cycles and iC321 in 10 MAGE cycles, corresponding to approximately 30 and 60 generations, respectively (one MAGE cycle amounts to ∼6 bacterial generations, on average^4^). This is on par with the ∼28 generations taken to construct the rationally-designed C.OPT—which showed poor growth characteristics in our studies—and is far fewer than the >1000 generations taken to generate C.M9 by adaptive evolution.

The findings from our study suggest that genome-scale codon reassignment may have smaller intrinsic fitness defects than is often assumed and that only a small subset of unintended mutations confer a dramatic impact on phenotype. The substantial fitness defects seen in recoded strains may instead be due to the exacerbation of dysregulated (as is the case for *ilvG*. Δ(A979-T980)) or suboptimal (as is the case for *prfB*.A246T) traits already present within ancestral strains. The mechanisms by which codon reassignment elicits this effect remain unclear, but our finding that reverting two K-12 lineage mutations leads to a substantial resolution of C321’s growth impairments suggests that recoding unmasks these metabolic idiosyncrasies. For this reason, we recommend that future recoding efforts be based off of high-fitness strains. The advantages (*i.e.* smaller genome size, absence of transposable genomic elements) and disadvantages (*i.e.* metabolic insufficiencies, physiological impairments) of organisms with minimal genomes^11,41^ should be carefully considered before using such strains as starting points for genome-scale engineering.

## Conclusion

Genomically recoded organisms hold promise for diverse biotechnological applications, but also exhibit fitness impairments that limit their utility. Here, we used targeted metabolic screening to inform the construction of engineered C321 derivatives with improved functional phenotypes. In addition to demonstrating superior growth kinetics in rich (**Supplementary data figure 5**) and minimal (**Figure 2E**) media, iC321 exhibits an EcNR2-like proteomic profile (**Figure 3C**), enhanced protein expression in minimal medium (**Figure 4B**), and high nsAA incorporation specificity in rich medium (**Figures 4D, F**). We anticipate that the strain will be useful for a wide variety of applications under a diverse range of growth conditions. However, iC321 does not outperform other recoded strains in all categories: C321 shows higher nsAA incorporation specificity in minimal medium than do iC321 and C321.M9 (**Figures 4E, G**), and C321.M9 grows faster than iC321 in minimal medium (**Figure 2E**). We suspect that the diverse metabolic and phenotypic characteristics of C321 derivatives will render certain strains more suitable for a particular application than others. iC321, given its improved growth kinetics and proteomic profile in minimal medium, may be particularly suitable for applications involving glucose metabolism. The favorable growth kinetics of C321.*prfB*^+^.*ilvG*^+^ with all seven C321 lineage reversions in rich medium may make it particularly suitable as a host strain for biofermentation. We recommend that as many strains as possible—both engineered and unoptimized—be tested when using genomically recoded organisms for any desired application, and that organisms constructed through future genomic recoding efforts undergo phenotype optimization to maximize their utility.

## Materials and methods

### Materials, strains and media

All strains engineered in this study are derived from *Escherichia coli* EcNR2 (Addgene ID 26931) and C321.ΔA (Addgene ID 48998). *E. coli* strains MG1655 and BL21 were obtained from the Coli Genetic Stock Center (CGSC# 6300 and 12504, respectively). C321.OPT and C321.M9 were obtained from the laboratory of Dr. George Church via Addgene (Addgene IDs 87359 and 98568, respectively). LB min media from AmericanBio (Canton, MA) was used for routine strain growth and cloning. M9 minimal medium was supplemented with 0.4% (w/v) glucose as a carbon source, 0.083 nM thiamine, and 5 nM biotin; additional supplements added to M9 for dropout experiments are detailed in **Supplementary table 2** and are based on concentrations used in Neidhardt rich defined medium^42^. 50 µg/mL carbenicillin was used to maintain cultures of EcNR2 and engineered derivatives, C321 and engineered derivatives, and C321.M9. 5 µg/mL gentamycin was used to maintain cultures of C321.OPT. MG1655 and BL21 were grown in antibiotic-free media. All cultures were grown at 34 °C.

### Serial passaging

Single colonies of C321 were picked from LB agar plates and inoculated into 3 mL LB or M9 supplemented with 50 µg/mL carbenicillin. Cultures were grown to saturation (*i.e.* overnight) and back-diluted 1:100 into 3 mL fresh media. This process was repeated daily for 10 days (for LB passaging) or 8 days (for M9 passaging). After passaging, single colonies were isolated by plating on LB agar and were analyzed via growth assays (detailed below).

### Plate-based growth assays

To determine kinetic growth parameters, we isolated single colonies of strains by plating cultures on LB medium containing appropriate antibiotics. Colonies (n = 3) were then inoculated into 3 mL LB and grown to an OD_600_ of 0.4-0.6. 1 mL of culture was washed by centrifugation (12,000 x *g*, 1 min) and resuspended in 1 mL M9 medium to remove residual nutrients from rich medium. Washed cultures were then diluted 1:100 into the wells of a flat-bottom 96-well bioassay plate containing 148.5 µL media and the appropriate antibiotics. Cultures were grown at 34°C in a Biotek Synergy HT microplate reader and OD_600_ was measured every 10 minutes.

### Calculation of growth parameters

Growth curves derived from 96-well microplate assays were analyzed in MATLAB (**Supplementary file 1**; also available as a GitHub repository (https://github.com/colinfromtherandomforest/cell-in-a-well). Doubling times were calculated by determining growth rate with the exponential phase of bacterial growth curves (*i.e.* when a log-transform of the growth curve is linear). The slope of the steepest portion of the log-transformed growth curve over 7 consecutive timepoints (*i.e.* one hour) is used to determine growth rate.

A common practice when calculating growth rates using this method is to blank all wells of a plate by subtracting the mean OD of several control wells containing media but no bacterial culture^20^. However, baseline OD_600_ values vary between wells on the same plate (variance ∼5% of mean), so the subtraction of a constant baseline from all plate wells affects doubling time calculations differently for each well. Furthermore, the subtraction of empty well OD values can lead to the calculation of artifactually low doubling times as the steepest portion of the log-transformed OD curve falls in a region within which the OD measured by the plate reader is nonlinear with respect to cell number (**Supplementary data figure 6**).

To avoid these artifacts, growth curves from all wells of a plate were adjusted to the same starting OD at the first timepoint. We set the appropriate starting OD for LB- and M9-containing wells separately by determining the threshold OD value below which the nonlinear effects of plate reader signal gain become significant, and setting a starting OD above this threshold (0.030 for LB and 0.020 for M9) (**Supplementary data figure 6**, dashed red lines). Doubling times are reported in minutes for experiments in which all strains were cultured within the same 96-well plate (*i.e.* **Figure 1B, 1C**). For experiments spanning multiple plates (*i.e.* **Figure 2E, 2F**), control wells containing EcNR2 and C321 were included in each plate to enable comparisons, and doubling times are reported as values normalized to those calculated for C321 in each plate. Raw and normalized growth parameters for these experiments are available in **Supplementary table 5**. OD_600_ values are reported as obtained by the plate reader and are not normalized to a 1cm path length.

### Strain engineering with targeted mutagenesis

We used MAGE^18,19^ to engineer all targeted genomic changes described in the study. Combinatorial genetic changes were implemented by performing three rounds of MAGE using pools of 90-mer single-stranded DNA oligonucleotides encoding the desired mutation (**Supplementary table 4**). Cultures were then plated on LB agar to isolate single bacterial colonies, which were screened for genotype using multiplex allele-specific colony PCR^18^. Additional rounds of MAGE were performed as needed to obtain desired genotypes. All genotypes were confirmed via Sanger sequencing.

### Cell culture for proteomics

Single colonies of strains analyzed in proteomics experiments were isolated by plating overnight on LB. Colonies (n = 3) were picked, inoculated into 3 mL M9 medium, and grown to mid-log phase (OD_600_ 0.4-0.6). Upon reaching mid-log, 1 mL of culture was transferred to 99 mL of pre-warmed M9 medium in a 250 mL culture flask. Flasks were incubated at 34 °C with shaking until cultures reached an OD_600_ of 0.5 to 0.6; because growth rates differ among strains, each flask was monitored for OD_600_ and removed from individually. Because C321 reaches lower maximum OD_600_ than other strains in minimal medium (**Figure 1A**), C321 cultures were removed at OD_600_ = 0.35. Cultures were concentrated into 1.5 mL microcentrifuge tubes and pellets were frozen at -80°C prior to proteomic work-up.

### Digestion of intact E. coli for shotgun proteomics

For cell lysis and protein digest, cell pellets were thawed on ice and 2 µL of cell pellet was transferred to a microcentrifuge tube containing 40 µL of lysis buffer (10 mM Tris-HCl pH 8.6, 10 mM DTT, 1 mM EDTA, and 0.5 % ALS). Cells were lysed by vortex for 30 seconds and disulfide bonds were reduced by incubating the reaction for 30 min. at 55°C. The reaction was briefly quenched on ice and 16 µL of a 60 mM IAA solution was added. Alkylation of cysteines proceeded for 30 min in the dark. Excess IAA was quenched with 14 µL of a 25 mM DTT solution and the sample was then diluted with 330 µL of 183 mM Tris-HCl buffer pH 8.0 supplemented with 2 mM CaCl_2_. Proteins were digested overnight using 12 μg sequencing grade trypsin. Following digestion, the reaction was then quenched with 12.5 µL of a 20% TFA solution, resulting in a sample pH ∼2. Remaining ALS reagent was cleaved for 15 min at room temperature. The sample (∼30 μg protein) was desalted by reverse phase clean-up using C18 MicroSpin columns (The Nest Group #SEM SS18V). Briefly, the column was conditioned twice with 400 µL 80% ACN, 0.1% TFA and twice with 400 µL 0.1% TFA by centrifugation. Samples were pelleted at 2000g for 1 minute before applying lysate to the column. Columns were then washed twice with 400 µL 0.1% TFA and eluted twice with 200 µL 80% ACN, 0.1% TFA. The desalted peptides were dried at room temperature in a rotary vacuum centrifuge and reconstituted in 30 µL 70% formic acid 0.1% TFA (3:8 v/v) for peptide quantitation by UV280. The sample was diluted to a final concentration of 0.2 μg/µL with 5 µL (1 μg) injected for LC-MS/MS analysis.

### Proteomics data acquisition and analysis

LC-MS/MS was performed using an ACQUITY UPLC M-Class (Waters) and Thermo Q Exactive Plus mass spectrometer. The analytical column employed was a 65-cm-long, 75-μm-internal-diameter PicoFrit column (New Objective) packed in-house to a length of 50 cm with 1.9 μm ReproSil-Pur 120 Å C18-AQ (Dr. Maisch) using methanol as the packing solvent. Peptide separation was achieved using mixtures of 0.1% formic acid in water (solvent A) and 0.1% formic acid in acetonitrile (solvent B) with a 90-min gradient 0/1, 2/7, 60/24, 65/48, 70/80, 75/80, 80/1, 90/1; (min/%B, linear ramping between steps). Gradient was performed with a flowrate of 250 nL/min. A single blank injection (5 µL 2% B) was performed between samples to eliminate peptide carryover on the analytical column. 100 fmol of trypsin-digested BSA or 100 ng trypsin-digested wildtype K-12 MG1655 *E. coli* proteins were run periodically between samples as quality control standards. The mass spectrometer was operated with the following parameters: (MS1) 70,000 resolution, 3e6 AGC target, 300–1,700 m/z scan range; (data dependent-MS2) 17,500 resolution, 1e6 AGC target, top 10 mode, 1.6 m/z isolation window, 27 normalized collision energy, 90s dynamic exclusion, unassigned and +1 charge exclusion. Data was searched using Maxquant version 1.6.10.43 with Deamidation (NQ) and Oxidation (M) as variable modifications and Carbamidomethyl (C) as a fixed modification with up to 3 missed cleavages, 5 AA minimum length, and 1% FDR against targeted libraries. *E. coli* proteome searches were run against a modified Uniprot *E. coli* database (taken on February 9, 2021), including full length IlvG. Proteome search results were analyzed with Perseus version 1.6.2.2 using a two-tailed Student’s t-test. If a protein was not detected in two of three replicate runs, it was designated as “not detected;” if the protein was not detected in one of three replicate runs, the non-detected sample was dropped from analysis. The MS proteomics data have been deposited to the ProteomeXchange Consortium via the PRIDE^43^ partner repository.

### Gene enrichment analysis

We performed Gene Ontology (GO) enrichment analysis on significant gene sets using the PANTHER classification system (www.pantherdb.org)^31^ using the *E. coli* all-gene reference list and the GO biological process annotation list. Enrichment analysis searches for GO terms that are over- and under-represented within input gene sets. The inputs for our analyses were the sets of all genes in EcNR2, C321.*prfB*^+^.*ilvG*^+^, and iC321 whose protein products showed significant differences in abundance compared to C321 (**Supplementary table 8**). Complete results of the enrichment analyses are given in **Supplementary table 9**.

### Plasmids for protein production and OTS activity assays

The pEVOL-pBpA plasmid, which encodes MjTyrRS/tRNA pair for incorporation of pBpA into proteins in *E. coli* in response to the amber codon (UAG), was obtained from the laboratory of Peter Schultz (Addgene #31190). pEvol-pBpA was maintained with 20 µg/mL chloramphenicol and induced using 0.2% w/v arabinose. ELP-0UAG-GFP, ELP-1UAG-GFP and ELP-3UAG-GFP fluorescence reporters^5^ were synthesized as gBlocks by IDT and inserted into plasmid backbones with a CloDF13 origin of replication and a spectinomycin resistance cassette using Gibson isothermal assembly (New England Biolabs, Boston, MA). Reporter plasmids were maintained using 95 µg/mL spectinomycin and induced using 30 ng/mL anhydrotetracycline (aTc).

### Endpoint protein production and OTS activity assays

ECNR2, C321, C321.M9, and iC321 cells were transformed with the following plasmid pairs: (1) pEvol-pBpA and ELP-0UAG-GFP, (2) pEvol-pBpA and ELP-1UAG-GFP, (3) pEvol-pBpA and ELP-3UAG-GFP. All transformed strains were grown at 34°C in LB medium supplemented with 20 µg/mL chloramphenicol and 95 µg/mL of spectinomycin. Wells of a 96-well plate were filled with 150 µL of LB medium supplemented with 20 µg/mL chloramphenicol and 95 µg/mL of spectinomycin. The wells were inoculated with colonies from each plasmid combination above (n = 3 biological replicates), and incubated at 34**°**C for 16 h with shaking. For endpoint protein production measurements, clear bottom wells of another 96-well plate were filled with 150 µL LB medium or M9 medium supplemented with 20 µg/mL chloramphenicol, 95 µg/mL of spectinomycin, and 30 ng/mL aTc to induce ELP-GFP expression. For measurement of OTS activity, clear bottom wells of a 96-well plate were filled with 150 µL of LB medium, or M9 medium, supplemented with 20 µg/mL chloramphenicol, 95 µg/mL of spectinomycin, 0.2% arabinose and 30 ng/mL aTc to induce OTS and ELP-GFP expression, respectively, and/or supplemented with 1mM *para*-benzoyl-L-phenylalanine (pBpa), where indicated. Cells from overnight growth were washed twice by centrifugation in 1X PBS to remove residual nutrients from rich medium, and 2 µL of culture was inoculated into the assay plates. Assay plates were incubated with linear shaking (731 cycles per min) for 16 h at 34°C on a Biotek Synergy H1 plate reader, with measurements of cell density (OD_600_) and GFP fluorescence (excitation at 485 nm and emission 528 nm with sensitivity setting at 70) taken at 10-minute intervals.

## Data availability

Whole-genome sequencing data for precursor and engineered C321 strains (C321.ΔA, C321.*prfB*^+^, C321.*ilvG*^+^, C321.*prfB*^+^.*ilvG*^+^, C321.*prfB*^+^.*ilvG*^+^*.gpp*^+^, C321.*prfB*^+^.*ilvG*^+^*.folA*^+^, C321.*prfB*^+^.*ilvG*^+^*.cyaA*^+^, iC321, C321.*prfB*^+^.*ilvG*^+^ with 7 C321-specific mutation reversions, C321.OPT, and C321.M9) have been deposited to the Sequence Read Archive under the temporary accession SUB10261001. Mass spectrometry and proteomics data have been deposited to the ProteomeXchange Consortium via the PRIDE^43^ partner repository with the identifier PXD030359.

## Acknowledgements

The authors would like to acknowledge the members of the Isaacs and Rinehart labs for their insightful comments on this study.

## Supplementary data figures

**Supplementary data figure 1.**
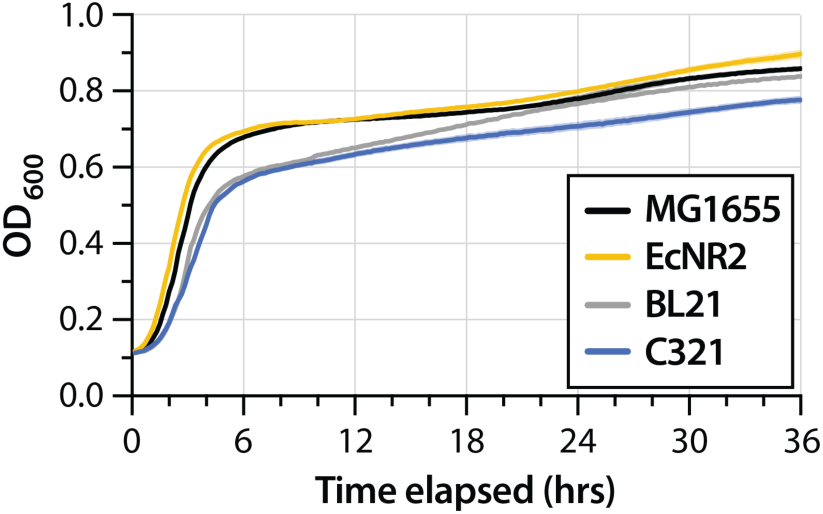
Growth curves of MG1655 (black), EcNR2 (yellow), non-K12 strain BL21 (grey), and C321 (blue) in LB rich media (n = 3 for all strains).

**Supplementary data figure 2.**
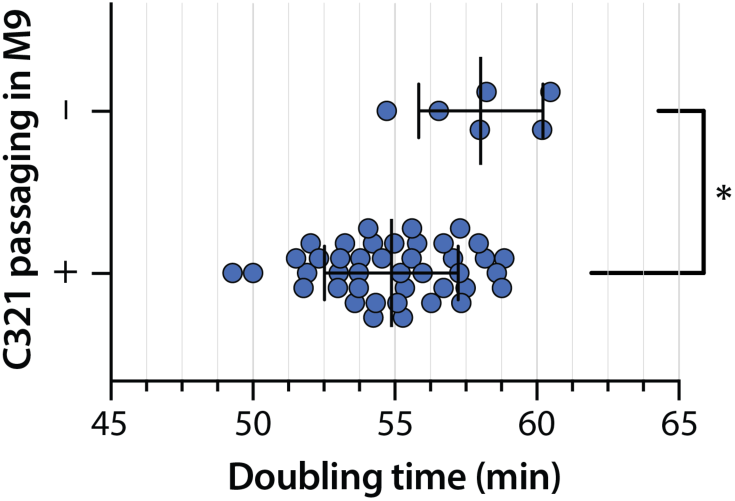
Doubling times of individual C321 clones with and without serial passaging in LB for ∼66 generations. Comparison of means by Welch’s *t*-test; p = 0.014.

**Supplementary data figure 3.**
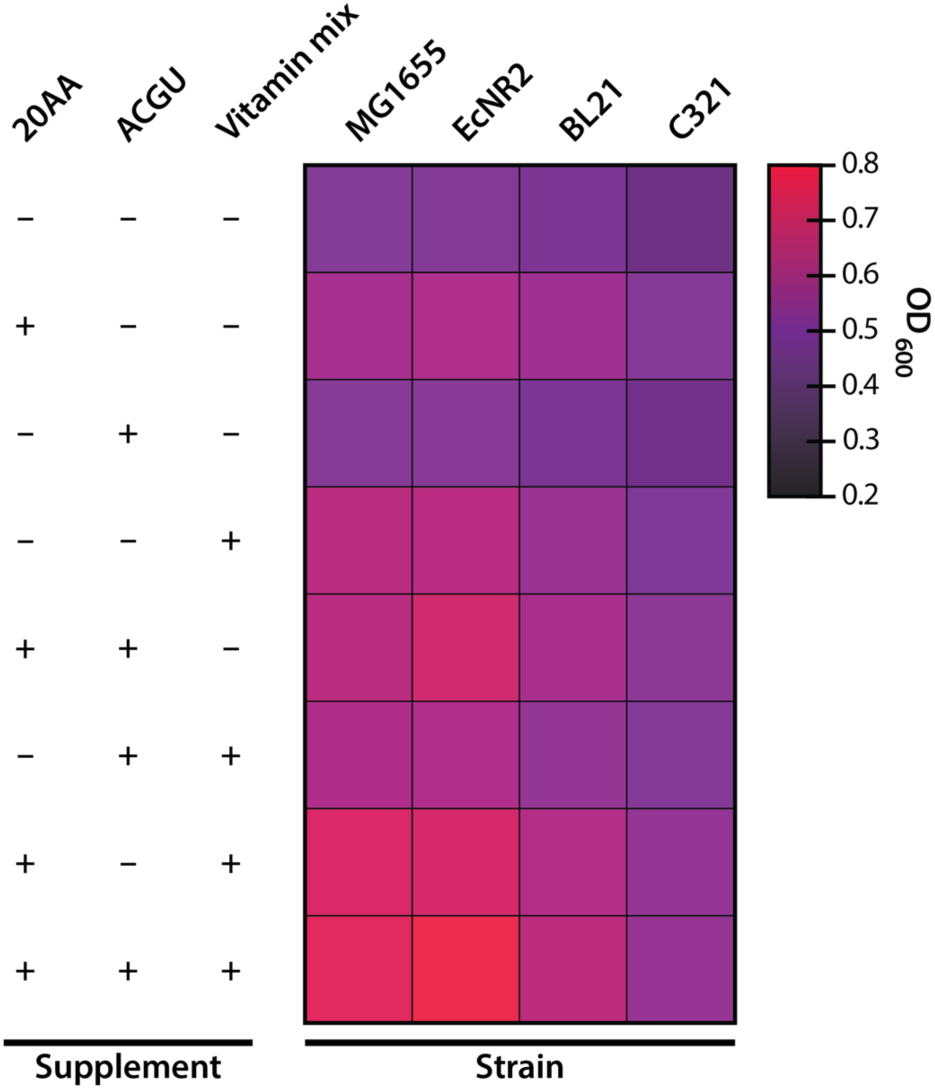
Maximum OD_600_ reached for C321 and related strains when grown in minimal media supplemented with 20 proteinogenic amino acids (20AA), 4 nucleobases (ACGU), and/or cofactors and trace metals (vitamin mix).

**Supplementary data figure 4.**
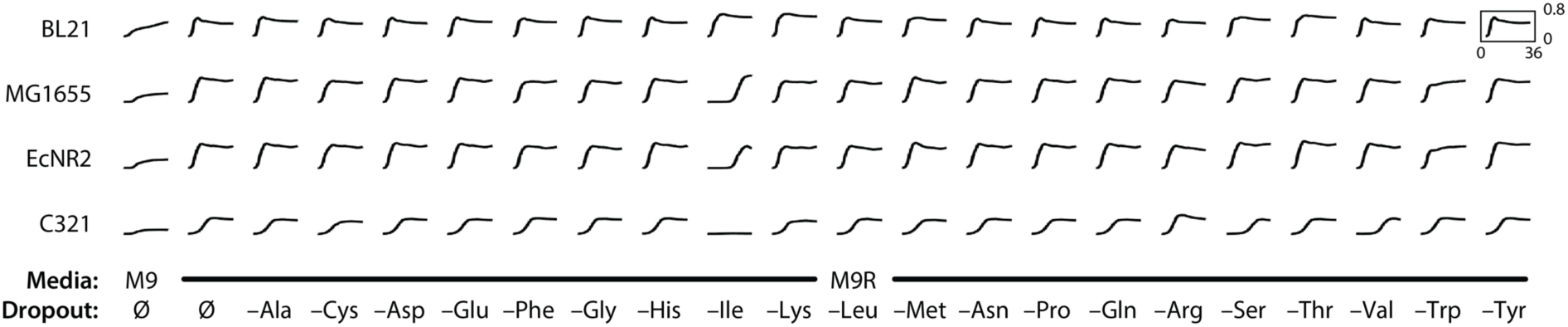
Growth curves for strains grown in M9, M9 medium supplemented with 20 amino acids (M9R), or M9R dropout medium lacking one of 20 amino acids. Each curve represents a 36-hour time course in a 96-well plate reader. All curves are scaled to a maximum OD_600_ of 0.8. The maximum OD_600_ reached in each condition is represented as a heatmap in main text figure 1D.

**Supplementary data figure 5.**
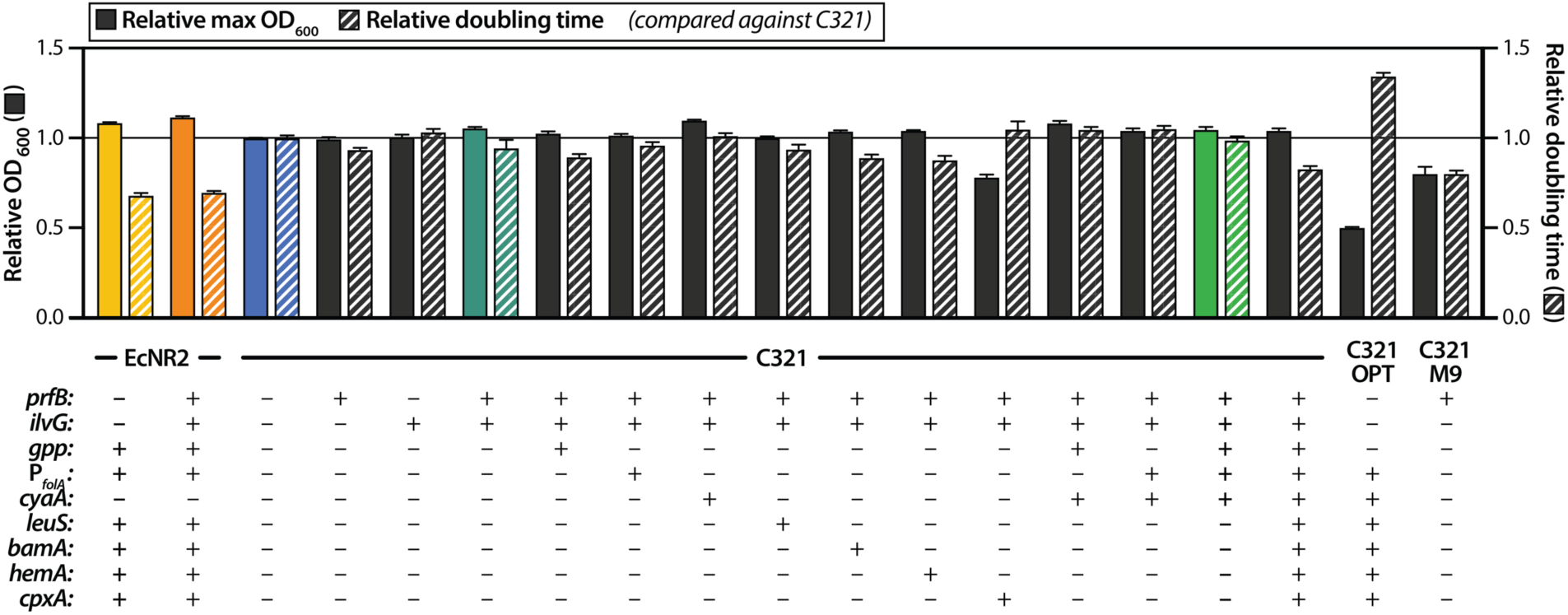
Maximum OD_600_ and doubling time relative to C321 for engineered C321 strains in LB. The solid horizontal line denotes the maximum OD_600_ and doubling time of C321. One-way ANOVA for comparisons between all strains is given in Supplementary table 6.

**Supplementary data figure 6.**
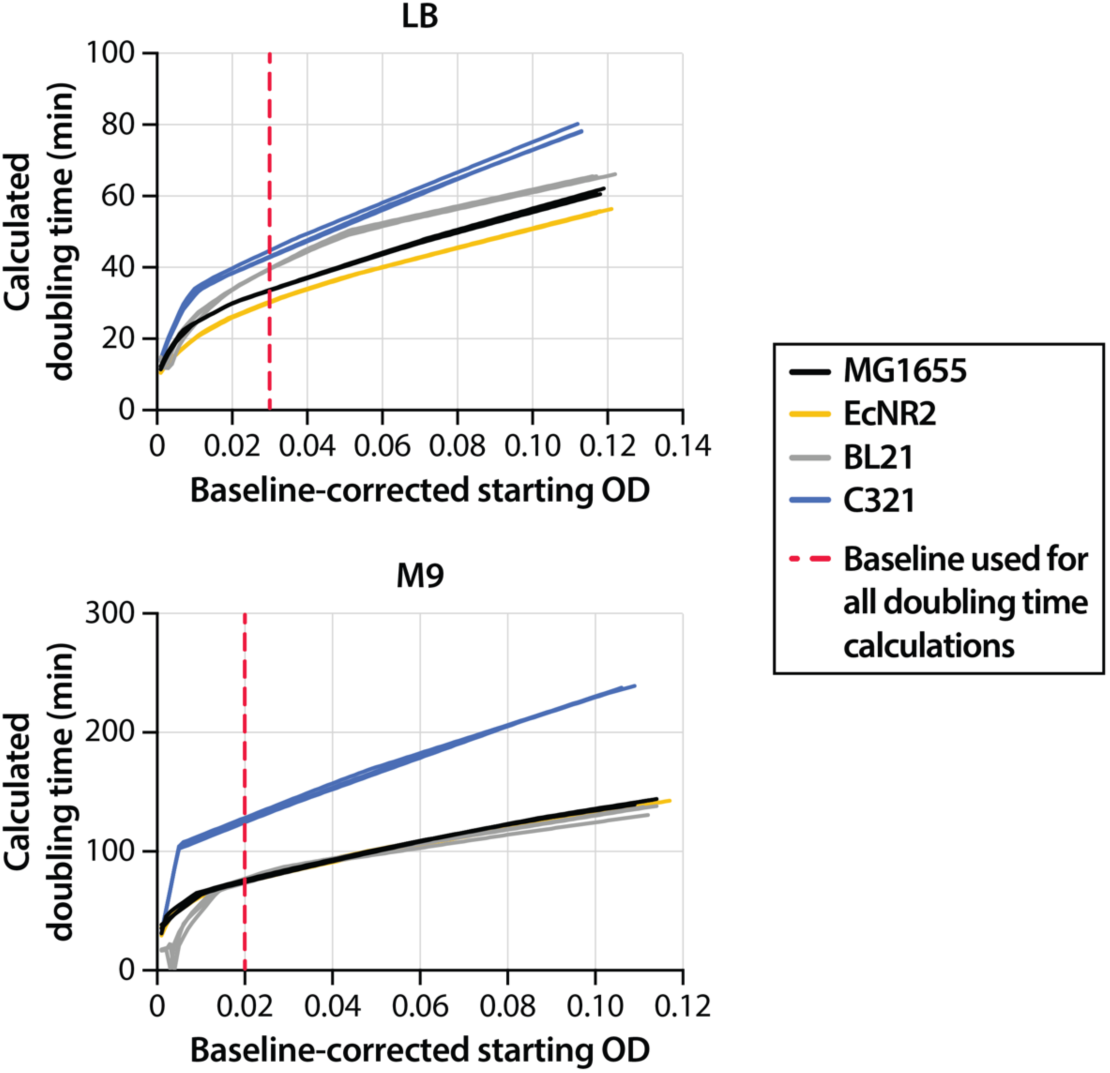
Effect of growth curve baseline correction on calculated doubling times. Doubling times were calculated for three replicates each of four strains in LB (top panel) and M9 (bottom panel) by shifting the growth curve to a defined starting OD value. Lines show calculated doubling times for each individual replicate. For all plate reader experiments reported in this study, growth curves were set to a baseline of 0.030 in LB and 0.020 in M9, represented by the dashed red lines.

## References

1 des Soye, B. J., Patel, J. R., Isaacs, F. J. & Jewett, M. C. Repurposing the translation apparatus for synthetic biology. Current Opinion in Chemical Biology 28, 83–90, doi:10.1016/j.cbpa.2015.06.008 (2015).

2 Ostrov, N. et al. Synthetic genomes with altered genetic codes. Current Opinion in Systems Biology 24, 32–40, doi:10.1016/j.coisb.2020.09.007 (2020).

3 Singh, T., Yadav, S. K., Vainstein, A. & Kumar, V. Genome recoding strategies to improve cellular properties: mechanisms and advances. Abiotech, 1–17, doi:10.1007/s42994-020-00030-1 (2020).

4 Lajoie, M. J. et al. Genomically recoded organisms expand biological functions. *Science (New York*, N.Y.) 342, 357–360, doi:10.1126/science.1241459 (2013).

5 Amiram, M. et al. Evolution of translation machinery in recoded bacteria enables multi-site incorporation of nonstandard amino acids. Nat Biotechnol 33, 1272–1279, doi:10.1038/nbt.3372 (2015).

6 Ma, N. J., Hemez, C. F., Barber, K. W., Rinehart, J. & Isaacs, F. J. Organisms with alternative genetic codes resolve unassigned codons via mistranslation and ribosomal rescue. eLife 7, e34878, doi:10.7554/eLife.34878 (2018).

7 Ma, N. J. & Isaacs, F. J. Genomic Recoding Broadly Obstructs the Propagation of Horizontally Transferred Genetic Elements. Cell Syst 3, 199–207, doi:10.1016/j.cels.2016.06.009 (2016).

8 Rovner, A. J. et al. Recoded organisms engineered to depend on synthetic amino acids. Nature 518, 89–93, doi:10.1038/nature14095 (2015).

9 Mandell, D. J. et al. Biocontainment of genetically modified organisms by synthetic protein design. Nature 518, 55–60, doi:10.1038/nature14121 (2015).

10 Ostrov, N. et al. Design, synthesis, and testing toward a 57-codon genome. *Science (New York*, N.Y.) 353, 819–822, doi:10.1126/science.aaf3639 (2016).

11 Fredens, J. et al. Total synthesis of Escherichia coli with a recoded genome. Nature 569, 514–518, doi:10.1038/s41586-019-1192-5 (2019).

12 Zhang, Y., Zhu, Y., Zhu, Y. & Li, Y. The importance of engineering physiological functionality into microbes. Trends Biotechnol 27, 664–672, doi:10.1016/j.tibtech.2009.08.006 (2009).

13 Lajoie, M. J. et al. Probing the limits of genetic recoding in essential genes. *Science (New York*, N.Y.) 342, 361–363, doi:10.1126/science.1241460 (2013).

14 Napolitano, M. G. et al. Emergent rules for codon choice elucidated by editing rare arginine codons in Escherichia coli. Proceedings of the National Academy of Sciences of the United States of America 113, E5588–5597, doi:10.1073/pnas.1605856113 (2016).

15 Lau, Y. H. et al. Large-scale recoding of a bacterial genome by iterative recombineering of synthetic DNA. Nucleic Acids Res 45, 6971–6980, doi:10.1093/nar/gkx415 (2017).

16 Richardson, S. M. et al. Design of a synthetic yeast genome. *Science (New York*, N.Y.) 355, 1040–1044, doi:10.1126/science.aaf4557 (2017).

17 Chen, Y. et al. Multiplex base editing to convert TAG into TAA codons in the human genome. 2021.2007.2013.452007 (2021).

18 Gallagher, R. R., Li, Z., Lewis, A. O. & Isaacs, F. J. Rapid editing and evolution of bacterial genomes using libraries of synthetic DNA. Nat Protoc 9, 2301–2316, doi:10.1038/nprot.2014.082 (2014).

19 Wang, H. H. et al. Programming cells by multiplex genome engineering and accelerated evolution. Nature 460, 894–898, doi:10.1038/nature08187 (2009).

20 Kuznetsov, G. et al. Optimizing complex phenotypes through model-guided multiplex genome engineering. Genome Biology 18, 100, doi:10.1186/s13059-017-1217-z (2017).

21 Wannier, T. M. et al. Adaptive evolution of genomically recoded Escherichia coli. Proceedings of the National Academy of Sciences 115, 3090–3095, doi:10.1073/pnas.1715530115 (2018).

22 Dragosits, M. & Mattanovich, D. Adaptive laboratory evolution -- principles and applications for biotechnology. Microb Cell Fact 12, 64, doi:10.1186/1475-2859-12-64 (2013).

23 Blattner, F. R. et al. The complete genome sequence of Escherichia coli K-12. *Science (New York*, N.Y.) 277, 1453–1462, doi:10.1126/science.277.5331.1453 (1997).

24 Andersen, D. C., Swartz, J., Ryll, T., Lin, N. & Snedecor, B. Metabolic oscillations in an E. coli fermentation. Biotechnology and Bioengineering 75, 212–218, doi:10.1002/bit.10018 (2001).

25 Uno, M., Ito, K. & Nakamura, Y. Functional specificity of amino acid at position 246 in the tRNA mimicry domain of bacterial release factor 2. Biochimie 78, 935–943, doi:10.1016/S0300-9084(97)86715-6 (1996).

26 Cashel, M. & Gallant, J. Two compounds implicated in the function of the RC gene of Escherichia coli. Nature 221, 838–841, doi:10.1038/221838a0 (1969).

27 Traxler, M. F. et al. The global, ppGpp-mediated stringent response to amino acid starvation in Escherichia coli. Molecular Microbiology 68, 1128–1148, doi:10.1111/j.1365-2958.2008.06229.x (2008).

28 Harms, E., Higgins, E., Chen, J. W. & Umbarger, H. E. Translational coupling between the ilvD and ilvA genes of Escherichia coli. J Bacteriol 170, 4798–4807, doi:10.1128/jb.170.10.4798-4807.1988 (1988).

29 Durfee, T., Hansen, A.-M., Zhi, H., Blattner, F. R. & Jin, D. J. Transcription profiling of the stringent response in Escherichia coli. J Bacteriol 190, 1084–1096, doi:10.1128/JB.01092-07 (2008).

30 Baccigalupi, L., Marasco, R., Ricca, E., De Felice, M. & Sacco, M. Control of ilvIH transcription during amino acid downshift in stringent and relaxed strains of Escherichia coli. FEMS Microbiol Lett 131, 95–98, doi:10.1111/j.1574-6968.1995.tb07760.x (1995).

31 Mi, H., Muruganujan, A., Casagrande, J. T. & Thomas, P. D. Large-scale gene function analysis with the PANTHER classification system. Nat Protoc 8, 1551–1566, doi:10.1038/nprot.2013.092 (2013).

32 Arranz-Gibert, P., Vanderschuren, K. & Isaacs, F. J. Next-generation genetic code expansion. Current Opinion in Chemical Biology 46, 203–211, doi:10.1016/j.cbpa.2018.07.020 (2018).

33 Chin, J. W., Martin, A. B., King, D. S., Wang, L. & Schultz, P. G. Addition of a photocrosslinking amino acid to the genetic code of Escherichiacoli. Proceedings of the National Academy of Sciences of the United States of America 99, 11020–11024, doi:10.1073/pnas.172226299 (2002).

34 Vyazmensky, M. et al. Interactions between large and small subunits of different acetohydroxyacid synthase isozymes of Escherichia coli. Biochemistry 48, 8731–8737, doi:10.1021/bi9009488 (2009).

35 Lee, H., Popodi, E., Tang, H. & Foster, P. L. Rate and molecular spectrum of spontaneous mutations in the bacterium Escherichia coli as determined by whole-genome sequencing. Proceedings of the National Academy of Sciences 109, E2774–E2783, doi:10.1073/pnas.1210309109 (2012).

36 Aakre, C. D. et al. Evolving new protein-protein interaction specificity through promiscuous intermediates. Cell 163, 594–606, doi:10.1016/j.cell.2015.09.055 (2015).

37 Kern, A. D. & Kondrashov, F. A. Mechanisms and convergence of compensatory evolution in mammalian mitochondrial tRNAs. Nat Genet 36, 1207–1212, doi:10.1038/ng1451 (2004).

38 Reynolds, M. G. Compensatory Evolution in Rifampin-Resistant Escherichia coli. Genetics 156, 1471–1481, doi:10.1093/genetics/156.4.1471 (2000).

39 Merker, M. et al. Compensatory evolution drives multidrug-resistant tuberculosis in Central Asia. eLife 7, e38200, doi:10.7554/eLife.38200 (2018).

40 Flemming, H.-C. & Wuertz, S. Bacteria and archaea on Earth and their abundance in biofilms. Nature Reviews Microbiology 17, 247–260, doi:10.1038/s41579-019-0158-9 (2019).

41 Pósfai, G. et al. Emergent properties of reduced-genome Escherichia coli. *Science (New York*, N.Y.) 312, 1044–1046, doi:10.1126/science.1126439 (2006).

42 Neidhardt, F. C., Bloch, P. L. & Smith, D. F. Culture Medium for Enterobacteria. J Bacteriol 119, 736–747 (1974).

43 Perez-Riverol, Y. et al. The PRIDE database and related tools and resources in 2019: improving support for quantification data. Nucleic Acids Res 47, D442–D450, doi:10.1093/nar/gky1106 (2019).

