## Supplementary File 1 for "Genomically recoded *Escherichia coli* with optimized functional phenotypes": README_iC321_SuppFile1.txt

This folder contains raw and processed data for growth curves reported in Hemez et al., "Genomically recoded Escherichia coli with optimized functional phenotypes."

File names: Each raw data file is named:

"iC321_[Main text figure]_[Supplementary figure]_[Media type]_[Experiment identifier]_[Experiment sub-identifier].xlsx"

The MAIN TEXT FIGURE and SUPPLEMENTARY TEXT FIGURE fields denote the manuscript figures in which the data contained within the file is reported. Other fields can be used by the authors to look up relevant notebook entries if needed. Each processed data file is named:

"OUTPUT_[Raw data file name].xls"

Processed data files are generated by running MATLAB scripts (supplementary file 2) that extract maximum OD600 values and doubling times from growth curves. Figures in which normalized doubling times are reported (i.e. Figure 1F, 2E, 2F) have additional OUTPUT files that contain formatted and normalized growth data. These additional files feature "_NORMALIZED" within their name. Data files that do not have an associated �OUTPUT_� file were not analyzed for doubling times (i.e. only raw growth curves or maximum OD values are reported).
