## Supplementary File 2 for "Genomically recoded *Escherichia coli* with optimized functional phenotypes": dataProcessing.m

```
%% Data processing function
% Colin Hemez
% GitHub: colinfromtherandomforest
% 2021

% Description: Takes an excel file of plate reader data and runs doubling
% time analysis on all wells.

% INPUTS
% WORKINGFILENAME: Excel file containing OD600 values for all samples to be
% analyzed. Time values sould be in the first column; names of samples
% should be in the first row.
% SSOD: Standard starting OD for all samples in the analysis, to be
% determined by the user. GROWTHCURVEANALYSIS will calculate doubling times
% both with and without a standard starting OD and will output both
% datasets into the results struct.

% OUTPUTS
% PROCESSEDDATA: Struct containing data processed from GROWTHCURVEANALYSIS
% (see GROWTHCURVEANALYSIS documentation for further information)
% DATAPROCESSING also writes an output file with all analyzed parameters
% for each well

function processedData = dataProcessing(workingFileName, ssOD)

workingTable = readtable([workingFileName,'.xlsx']);
workingData = table2array(workingTable(:,(2:end)));
writematrix(workingData,[workingFileName,'_RawCurves.xlsx'],  'FileType', 'spreadsheet');
processedData = growthCurveAnalysis([workingFileName,'_RawCurves.xlsx'],...
                               'standardstartod', ssOD,...
                               'summaryplotter', 'plotoff',...
                               'outputfilename', [workingFileName,'_Processed']);
processedFile = readtable([workingFileName,'_Processed.xls']);
strainID = workingTable.Properties.VariableNames(2:end)';
strainTable = table(strainID);
output = [strainTable processedFile];
writetable(output, ['OUTPUT_', workingFileName,'_Processed'], 'FileType', 'spreadsheet');

end
```
