## Supplementary File 2 for "Genomically recoded *Escherichia coli* with optimized functional phenotypes": growthCurveAnalysis.m

```
%% Plate reader growth curve analysis
% Colin Hemez
% GitHub: colinfromtherandomforest
% 2021

%%%%%%%%%%%%%%%%%%%%%%%%%%%%%%%%%%%%%%%%%%%%%%%%%%%%%%%%%%%%%%%%%%%%%%%%%%%
% REQUIRED INPUT                                                          %
% INPUTFILE: The file name for the raw OD values.                         %
%            Supported file types: .csv, .xls, .xlsx                      %
%%%%%%%%%%%%%%%%%%%%%%%%%%%%%%%%%%%%%%%%%%%%%%%%%%%%%%%%%%%%%%%%%%%%%%%%%%%

%%%%%%%%%%%%%%%%%%%%%%%%%%%%%%%%%%%%%%%%%%%%%%%%%%%%%%%%%%%%%%%%%%%%%%%%%%%
% OPTIONAL INPUTS                                                         %
% TIMEINTERVAL: Time interval (in minutes) between each OD measurement.   %
%               Default is 10 min.                                        %
% FIRSTTIMEPOINT: Time elapsed (in minutes) at which first OD measurement %
%                 was taken. Default is 10 min                            %
% DTPLOTTER: Specify whether to show figures from doubling time           %
%            calculation function. 'ploton' or 'plotoff'. Default is      %
%            'plotoff'.                                                   %
% STANDARDSTARTOD: Define the starting OD for standardized doubling time  %
%                  calculations. Default is 0.09.                         %
% BASELINESUBTRACT: Define a baseline OD value to subtract from all wells %
%                   in the analysis. If a vaseline subtraction value is   %
%                   specified, STANDARDSTARTOD will be ignored.           %
% CONTROLSTRAININDEX: Specify the index of the 'control' strain against   %
%                     which other strains are compared. Index is the      %
%                     column number of the 'control' strain in the input  %
%                     data file. Input must be an integer value.          %
%                     Default is 1.                                       %
% DTTIMEPOINTS: Specifies the number of timepoints to use in doubling     %
%               time calculations when checking for log linearity. Input  %
%               must be an odd integer. Default is 7.                     %
% SUMMARYPLOTTER: Specify whether to show summary plots of data.          %
%                 'ploton' or 'plotoff'. Default is 'plotoff'.            %
% OUTPUTFILENAME: Name of the output file. Must be entered as a string of %
%                 the form 'OUTPUTFILENAME.xlsx'. Of not specified,       %
%                 program names the output file using the current time.   %
% HIGHLIGHTSAMPLE: Specify the index of a strain to be highlighted in     %
%                  summary plots. Any number of strains can be specified. %
%                  To specify multiple strains, enter indices as a        %
%                  vector. Default is [].                                 %
% PLATESUBSET: Define the subset of samples to analyze. Input can be a    %
%              vector of integers that denote the indices of strains to   %
%              be analyzed. Input can also be a string that specifies a   %
%              predefined plate subset:                                   %
%              STRING       WELLS                                         %
%              tophalf      A1-D12 (top 4 rows)                           %
%              bottomhalf   E1-H12 (bottom 4 rows)                        %
%              lefthalf     A1-A6, B1-B6, ... , G1-G6, H1-H6              %
%              righthalf    A7-A12, B7-B12, ... , G7-G12, H7-H12          %
% TIMESUBSET: Define a subset of timepoints to analyze. Input must be a   %
%             two-value vector of the form [tmin tmax]. Values for tmin   %
%             (earliest timepoint) and tmax (latest timepoint) are in     %
%             minutes.                                                    %
%                                                                         %
% All optional inputs are entered as name-value pairs                     %
%%%%%%%%%%%%%%%%%%%%%%%%%%%%%%%%%%%%%%%%%%%%%%%%%%%%%%%%%%%%%%%%%%%%%%%%%%%

%%%%%%%%%%%%%%%%%%%%%%%%%%%%%%%%%%%%%%%%%%%%%%%%%%%%%%%%%%%%%%%%%%%%%%%%%%%
% OUTPUTS                                                                 %
% PROCESSEDSTRUCT: Processed data, returned as a struct.                  %
% RAW: Input OD vaues pulled from INPUTFILE, returned as a matrix. This   %
%      output returns all of the data contained in INPUTFILE, and not     %
%      just the subset for analysis specified by PLATESUBSET.             %
% PROCESSEDTABLE: Processed data, returned as a table.                    %
% TIMEVALS: Array of time values (in minutes) at which OD measurements    %
%           were taken. This output returns all of the time values for    %
%           all timepoints defined in RAW, and not just the subset of     %
%           time values specified by TIMESUBSET.                          %
%%%%%%%%%%%%%%%%%%%%%%%%%%%%%%%%%%%%%%%%%%%%%%%%%%%%%%%%%%%%%%%%%%%%%%%%%%%

%%%%%%%%%%%%%%%%%%%%%%%%%%%%%%%%%%%%%%%%%%%%%%%%%%%%%%%%%%%%%%%%%%%%%%%%%%%
% DEPENDENCIES                                                            %
% Function lnDoublingTime() must be in the same directory                 %
%%%%%%%%%%%%%%%%%%%%%%%%%%%%%%%%%%%%%%%%%%%%%%%%%%%%%%%%%%%%%%%%%%%%%%%%%%%

function [processedstruct, raw, processedtable, timevals] = ...
         growthCurveAnalysis(inputfile, varargin)

tic
disp('   ')
disp('growthCurveAnalysis()...')
disp('   ')

% Define default inputs
defaultTimeInt = 10;
defaultFirstTimePoint = 10;
defaultDTPlotter = 0;
defaultStandardStartOD = 0.09;
defaultBaselineSubtraction = 0;
defaultControlStrainIndex = 1;
defaultDTTimePoints = 7;
defaultSummaryPlotter = 'ploton';
defaultHighlightSample = [];
defaultPlateSubset = [];
defaultTimeSubset = [-inf inf];

% Define default output file name
filenoext = inputfile;
while ~strcmp(filenoext(end), '.')
    filenoext = filenoext(1:(end-1));
end
filenoext = filenoext(1:(end-1));
currtime = datetime('now');
currtime = datestr(currtime);
defaultOutputFilename = [filenoext '_Processed_' currtime '.xlsx'];

% Parse inputs
p = inputParser;
addRequired(p, 'inputfile');
addOptional(p, 'timeinterval', defaultTimeInt)
addOptional(p, 'firsttimepoint', defaultFirstTimePoint)
addOptional(p, 'dtplotter', defaultDTPlotter)
addOptional(p, 'standardstartod', defaultStandardStartOD)
addOptional(p, 'baselinesubtract', defaultBaselineSubtraction)
addOptional(p, 'controlstrainindex', defaultControlStrainIndex)
addOptional(p, 'dttimepoints', defaultDTTimePoints)
addOptional(p, 'summaryplotter', defaultSummaryPlotter)
addOptional(p, 'outputfilename', defaultOutputFilename)
addOptional(p, 'highlightsample', defaultHighlightSample)
addOptional(p, 'platesubset', defaultPlateSubset)
addOptional(p, 'timesubset', defaultTimeSubset)
parse(p, inputfile, varargin{:});

% Unpack parameters from parsing results
timeint = p.Results.timeinterval;
firsttimepoint = p.Results.firsttimepoint;
dtplotter = p.Results.dtplotter;
standardstartod = p.Results.standardstartod;
baselinesubtract = p.Results.baselinesubtract;
controlstrainindex = p.Results.controlstrainindex;
dttimepoints = p.Results.dttimepoints;
summaryplotter = p.Results.summaryplotter;
outputfilename = p.Results.outputfilename;
highlightsample = p.Results.highlightsample;
platesubset = p.Results.platesubset;
timesubset = p.Results.timesubset;

% Determine if a plate subset is specified
if ~isempty(platesubset)
    if ischar(platesubset)
        % Preset plate subset schemes
        platesubset = lower(platesubset);
        if strcmp(platesubset, 'tophalf')
            subsetind = 1:48;
        elseif strcmp(platesubset, 'bottomhalf')
            subsetind = 49:96;
        elseif strcmp(platesubset, 'lefthalf')
            subsetind = [1:6 13:18 25:30 37:42 49:54 61:66 73:78 85:90];
        elseif strcmp(platesubset, 'righthalf')
            subsetind = [1:6 13:18 25:30 37:42 49:54 61:66 73:78 85:90]+6;
        else
            error('Invalid option specified for PLATESUBSET')
        end
        
        internalind = find(subsetind == controlstrainindex);

        % Check to make sure that the control sample is within subset range
        if sum(controlstrainindex == subsetind) == 0
            warning(['Control sample index is not within plate ', ...
                     'subset range. Setting control to index ', ...
                     num2str(min(subsetind)), '.'])
            controlstrainindex = min(subsetind);
            internalind = find(subsetind == controlstrainindex);
        end
        
    else
        % Custom-defined plate subset scheme
        subsetind = platesubset;
        internalind = find(subsetind == controlstrainindex);
        
        % Check to make sure that a control strain exists
        if isempty(internalind)
            warning(['Control sample index not specified and default ', ...
                     'is out of range. Setting control to index ',...
                     num2str(min(subsetind)), '.'])
            controlstrainindex = min(subsetind);
            internalind = find(subsetind == controlstrainindex);
        end
    end
    
    % Find internal indices of highlight samples
    if ~isempty(highlightsample)
        S = zeros(1, length(subsetind));
        for i = 1:length(highlightsample)
            Hi = highlightsample(i);
            Si = (subsetind == Hi);
            S = S + Si;
        end
        highlightinternalind = find(S);
    end
    
end

% Determine whether to subtract a baseline or standardize the starting OD
if baselinesubtract ~= 0
    baselineflag = true;
    disp(['Subtracting a constant baseline (',...
          num2str(baselinesubtract),...
          ') from all growth curves'])
else
    baselineflag = false;
    disp(['Setting all growth curves to a standard starting OD (',...
          num2str(standardstartod),...
          ')'])
end

% Load data file and calculate dimensions
rawdatafull = readmatrix(inputfile);
if ~isempty(platesubset)
    rawdata = rawdatafull(:, subsetind);
else
    rawdata = rawdatafull;
end
dim = size(rawdata);
nsamples = dim(2);
if isempty(platesubset)
    subsetind = 1:nsamples;
    if ~isempty(highlightsample)
        S = zeros(1, nsamples);
        for i = 1:length(highlightsample)
            Hi = highlightsample(i);
            Si = (subsetind == Hi);
            S = S + Si;
        end
        highlightinternalind = find(S);
    end
end
ntimepoints = dim(1);

% Make the time vector
timeind = 0:1:(ntimepoints-1);
timefull = timeind .* timeint + firsttimepoint;
timefull = timefull';
time = timefull;

if sum(timesubset == defaultTimeSubset) < 2
    % Constrain time vector by parameters specified in timesubset
    timelogical = (timefull >= timesubset(1) & timefull <= timesubset(2));
    time = timefull(timelogical);
    
    % Constrain OD data matrix by parameters specified in timesubset
    datasubtime = zeros(length(time), nsamples);
    for i = 1:nsamples
        datasubi = rawdata(:,i);
        datasubtimei = datasubi(timelogical);
        datasubtime(:,i) = datasubtimei;
    end
    
    rawdata = datasubtime;
    
end

% Determine whether or not to plot individual doubling time calculations
if dtplotter == 0
    dtplotfcn = 'plotoff';
elseif strcmp(dtplotter, 'ploton')
    dtplotfcn = 'ploton';
else
    error('Invalid input for argument DTPLOTTER')
end

%%%%%%%%%%%%%%%%%%%%%%%%%%%%%%%%%%%%%%%%%
%%%%%%%%%%%%%%%%%%%%%%%%%%%%%%%%%%%%%%%%%%%%
%%%%%%%%%%%%%%%%%%%%%%%%%%%%%%%%%%%%%%%%%%%%%%%
% Determine maximum OD values from the raw data
maxOD = max(rawdata);
maxOD = maxOD';
%%%%%%%%%%%%%%%%%%%%%%%%%%%%%%%%%%%%%%%%%%%%%%%
%%%%%%%%%%%%%%%%%%%%%%%%%%%%%%%%%%%%%%%%%%%%
%%%%%%%%%%%%%%%%%%%%%%%%%%%%%%%%%%%%%%%%%

% Generate OD matrix with standardized starting value
datastandardstart = zeros(length(time), nsamples);

for i = 1:nsamples
    curvei = rawdata(:,i);
    if baselineflag
        curvei = curvei - baselinesubtract;
    else
        curvei = curvei - min(curvei) + standardstartod;
    end
    datastandardstart(:,i) = curvei;
end

%%%%%%%%%%%%%%%%%%%%%%%%%%%%%%%%%%%%%%%%%
%%%%%%%%%%%%%%%%%%%%%%%%%%%%%%%%%%%%%%%%%%%%
%%%%%%%%%%%%%%%%%%%%%%%%%%%%%%%%%%%%%%%%%%%%%%%
% Calculate the doubling times for the raw data
rawDT = zeros(nsamples, 1);
rawRsq = zeros(nsamples, 1);
rawTimeMedian = zeros(nsamples, 1);

for i = 1:nsamples
    curvei = rawdata(:,i);
    [DTi, Ri, Mi] = lnDoublingTime(curvei, time, dttimepoints, dtplotfcn);
    rawDT(i) = DTi;
    rawRsq(i) = Ri;
    rawTimeMedian(i) = Mi;
end
%%%%%%%%%%%%%%%%%%%%%%%%%%%%%%%%%%%%%%%%%%%%%%%
%%%%%%%%%%%%%%%%%%%%%%%%%%%%%%%%%%%%%%%%%%%%
%%%%%%%%%%%%%%%%%%%%%%%%%%%%%%%%%%%%%%%%%


%%%%%%%%%%%%%%%%%%%%%%%%%%%%%%%%%%%%%%%%%
%%%%%%%%%%%%%%%%%%%%%%%%%%%%%%%%%%%%%%%%%%%%
%%%%%%%%%%%%%%%%%%%%%%%%%%%%%%%%%%%%%%%%%%%%%%%
% Calculate the doubling times for the OD standardized data
startnormDT = zeros(nsamples, 1);
startnormRsq = zeros(nsamples, 1);
startnormTimeMedian = zeros(nsamples, 1);

for i = 1:nsamples
    curvei = datastandardstart(:,i);
    [DTi, Ri, Mi] = lnDoublingTime(curvei, time, dttimepoints);
    startnormDT(i) = DTi;
    startnormRsq(i) = Ri;
    startnormTimeMedian(i) = Mi;
end

% Calculate the relative doubling time factors
if ~isempty(platesubset)
    relDT = startnormDT ./ startnormDT(internalind);
else
    relDT = startnormDT ./ startnormDT(controlstrainindex);
end

%%%%%%%%%%%%%%%%%%%%%%%%%%%%%%%%%%%%%%%%%%%%%%%
%%%%%%%%%%%%%%%%%%%%%%%%%%%%%%%%%%%%%%%%%%%%
%%%%%%%%%%%%%%%%%%%%%%%%%%%%%%%%%%%%%%%%%


%%%%%%%%%%%%%%%%%%%%%%%%%%%%%%%%%%%%%%%%%
%%%%%%%%%%%%%%%%%%%%%%%%%%%%%%%%%%%%%%%%%%%%
%%%%%%%%%%%%%%%%%%%%%%%%%%%%%%%%%%%%%%%%%%%%%%%
% Plot summary data
if strcmp(summaryplotter, 'ploton')
    % Plot summary data
    
    % Plot raw starting OD values for samples with mean and uncertainty
    figure
    if ~isempty(platesubset)
        startingODs = rawdata(1,:);
        plot(controlstrainindex, startingODs(internalind), 'or',...
         'markerfacecolor', 'r', 'markersize', 8)
        hold on
        plot(subsetind, startingODs, 'ok',...
             'markerfacecolor', 'k', 'markersize', 8)
        plot(controlstrainindex, startingODs(internalind), 'or',...
             'markerfacecolor', 'r', 'markersize', 8)
        minind = min(subsetind);
        maxind = max(subsetind);
        M = median(startingODs);
        P10 = prctile(startingODs, 10);
        P90 = prctile(startingODs, 90);
        plot([minind maxind], [M M], '-k', 'linewidth', 2)
        plot([minind maxind], [P10 P10], '--k', 'linewidth', 2)
        plot([minind maxind], [P90 P90], '--k', 'linewidth', 2)
        xlim([(minind-1) (maxind+1)])
    else
        startingODs = rawdata(1,:);
        plot(controlstrainindex, startingODs(controlstrainindex), 'or',...
         'markerfacecolor', 'r', 'markersize', 8)
        hold on
        plot(1:nsamples, startingODs, 'ok',...
             'markerfacecolor', 'k', 'markersize', 8)
        plot(controlstrainindex, startingODs(controlstrainindex), 'or',...
             'markerfacecolor', 'r', 'markersize', 8)
        minind = min(1:nsamples);
        maxind = max(1:nsamples);
        M = median(startingODs);
        P10 = prctile(startingODs, 10);
        P90 = prctile(startingODs, 90);
        plot([minind maxind], [M M], '-k', 'linewidth', 2)
        plot([minind maxind], [P10 P10], '--k', 'linewidth', 2)
        plot([minind maxind], [P90 P90], '--k', 'linewidth', 2)
        xlim([(minind-1) (maxind+1)])
    end
    
    if ~isempty(highlightsample)
        plot(highlightsample, startingODs(highlightinternalind),...
             'o', 'markerfacecolor', '#EDB120',...
             'markeredgecolor', '#EDB120', 'markersize', 8)
    end
    
    grid on
    xlabel('Strain index')
    ylabel('Starting OD_{600}')
    title('Starting ODs for all samples')
    legend('Control strain', 'Other strains', 'location', 'northwest')
    
    % Plot growth curves for all samples
    figure
    
    if ~isempty(platesubset)
        plot(time./60, rawdata(:,internalind), '-r', 'linewidth', 4)
        hold on
        plot(time./60, rawdata, '-k', 'linewidth', 1)
        plot(time./60, rawdata(:,internalind), '-r', 'linewidth', 4)
    else
        plot(time./60, rawdata(:,controlstrainindex), '-r', 'linewidth', 4)
        hold on
        plot(time./60, rawdata, '-k', 'linewidth', 1)
        plot(time./60, rawdata(:,controlstrainindex), '-r', 'linewidth', 4)
    end
    
    if ~isempty(highlightsample)
        plot(time./60, rawdata(:,highlightinternalind), '-',...
             'linewidth', 4, 'color', '#EDB120')
    end
    
    grid on
    xlabel('Time (hrs)')
    ylabel('OD_{600}')
    title('Gowth curves for all samples')
    legend('Control strain', 'Other strains', 'location', 'northwest')
    
    % Dot plot the max OD and doubling time for all samples
    figure
    
    if ~isempty(platesubset)
        plot(maxOD(internalind), rawDT(internalind), 'or',...
         'markerfacecolor', 'r', 'markersize', 8)
        hold on
        plot(maxOD, rawDT, 'ok',...
             'markerfacecolor', 'k', 'markersize', 8)
        plot(maxOD(internalind), rawDT(internalind), 'or',...
             'markerfacecolor', 'r', 'markersize', 8)
    else
        plot(maxOD(controlstrainindex), rawDT(controlstrainindex), 'or',...
         'markerfacecolor', 'r', 'markersize', 8)
        hold on
        plot(maxOD, rawDT, 'ok',...
             'markerfacecolor', 'k', 'markersize', 8)
        plot(maxOD(controlstrainindex), rawDT(controlstrainindex), 'or',...
             'markerfacecolor', 'r', 'markersize', 8)
    end
    
    if ~isempty(highlightsample)
        plot(maxOD(highlightinternalind), rawDT(highlightinternalind),...
             'o', 'markerfacecolor', '#EDB120',...
             'markeredgecolor', '#EDB120', 'markersize', 8)
    end
    
    grid on
    xlabel('Maximum OD_{600}')
    ylabel('Doubling Time (min)')
    title('Maximum OD and doubling time for all samples')
    legend('Control strain', 'Other strains', 'location', 'northwest')
    
    % Plot histograms of max OD, raw DT, and OD-standardized relative DT
    figure
    
    % Maximum OD
    subplot(1,3,1)
    H = histogram(maxOD, 'facecolor', [0.5 0.5 0.5]);
    hold on
    
    if ~isempty(platesubset)
        plot([maxOD(internalind) maxOD(internalind)],...
             [0 max(H.Values)], '--r', 'linewidth', 3)
    else
        plot([maxOD(controlstrainindex) maxOD(controlstrainindex)],...
             [0 max(H.Values)], '--r', 'linewidth', 3)
    end
    
    if ~isempty(highlightsample)
        for i = 1:length(highlightsample)
            plot([maxOD(highlightinternalind(i))...
                  maxOD(highlightinternalind(i))],...
                 [0 max(H.Values)], '-', 'linewidth', 3,...
                 'color', '#EDB120')
        end
    end
    
    grid on
    xlabel('Maximum OD_{600}')
    ylabel('Frequency')
    title('Maximum OD distributions')
    legend('Other strains', 'Control strain', 'location', 'northwest')
    
    % Raw DT
    subplot(1,3,2)
    H = histogram(rawDT, 'facecolor', [0.5 0.5 0.5]);
    hold on
    
    if ~isempty(platesubset)
        plot([rawDT(internalind) rawDT(internalind)],...
             [0 max(H.Values)], '--r', 'linewidth', 3)
    else
        plot([rawDT(controlstrainindex) rawDT(controlstrainindex)],...
             [0 max(H.Values)], '--r', 'linewidth', 3)
    end
    
    if ~isempty(highlightsample)
        for i = 1:length(highlightsample)
            plot([rawDT(highlightinternalind(i))...
                  rawDT(highlightinternalind(i))],...
                 [0 max(H.Values)], '-', 'linewidth', 3,...
                 'color', '#EDB120')
        end
    end
    
    grid on
    xlabel('Doubling Time (min)')
    ylabel('Frequency')
    title('Raw doubling time distribution')
    legend('Other strains', 'Control strain', 'location', 'northwest')
    
    % Relative DT factors
    subplot(1,3,3)
    H = histogram(relDT, 'facecolor', [0.5 0.5 0.5]);
    hold on
    
    if ~isempty(platesubset)
        plot([relDT(internalind) relDT(internalind)],...
             [0 max(H.Values)], '--r', 'linewidth', 3)
    else
        plot([relDT(controlstrainindex) relDT(controlstrainindex)],...
             [0 max(H.Values)], '--r', 'linewidth', 3)
    end

    if ~isempty(highlightsample)
        for i = 1:length(highlightsample)
            plot([relDT(highlightinternalind(i))...
                  relDT(highlightinternalind(i))],...
                 [0 max(H.Values)], '-', 'linewidth', 3,...
                 'color', '#EDB120')
        end
    end
    
    grid on
    xlabel('Relative doubling time factor')
    ylabel('Frequency')
    title('Relative doubling time factor distribution')
    legend('Other strains', 'Control strain', 'location', 'northwest')
    
elseif strcmp(summaryplotter, 'plotoff')
    % Do not plot summary data
    disp('   ')
    disp('Summary plot generation suppressed')
    
else
    % Input not recognized
    disp('   ')
    error('Invalid input for argument SUMMARYPLOTTER')
    
end
%%%%%%%%%%%%%%%%%%%%%%%%%%%%%%%%%%%%%%%%%%%%%%%
%%%%%%%%%%%%%%%%%%%%%%%%%%%%%%%%%%%%%%%%%%%%
%%%%%%%%%%%%%%%%%%%%%%%%%%%%%%%%%%%%%%%%%


%%%%%%%%%%%%%%%%%%%%%%%%%%%%%%%%%%%%%%%%%
%%%%%%%%%%%%%%%%%%%%%%%%%%%%%%%%%%%%%%%%%%%%
%%%%%%%%%%%%%%%%%%%%%%%%%%%%%%%%%%%%%%%%%%%%%%%
% Package all output data into a matrix
processedmat = [subsetind'...
                maxOD...
                rawDT...
                rawRsq...
                rawTimeMedian...
                startnormDT...
                startnormRsq...
                startnormTimeMedian...
                relDT];
processedmatdim = size(processedmat);

% Prepare results table
variableNames = {'Strain_Index',...
                 'Max_OD',...
                 'DT_Raw',...
                 'DT_Raw_Rsq',...
                 'DT_Raw_Time_Median',...
                 'DT_Standardized',...
                 'DT_Standardized_Rsq',...
                 'DT_Standardized_Time_Median',...
                 'DT_Relative'};
variableTypes = {'double',...
                 'double',...
                 'double',...
                 'double',...
                 'double',...
                 'double',...
                 'double',...
                 'double',...
                 'double'};
T = table('Size', [nsamples length(variableNames)],...
          'VariableTypes', variableTypes,...
          'VariableNames', variableNames);
for i = 1:processedmatdim(2)
    T.(variableNames{i})(1:nsamples) = processedmat(:,i);
end
%%%%%%%%%%%%%%%%%%%%%%%%%%%%%%%%%%%%%%%%%%%%%%%
%%%%%%%%%%%%%%%%%%%%%%%%%%%%%%%%%%%%%%%%%%%%
%%%%%%%%%%%%%%%%%%%%%%%%%%%%%%%%%%%%%%%%%

% Write the table as an excel file
writetable(T, outputfilename, 'FileType', 'spreadsheet');

% Return the function outputs
raw = rawdatafull;
processedtable = T;
for i = 1:processedmatdim(2)
    processedstruct.(variableNames{i}) = processedmat(:,i);
end
timevals = timefull;

% Display some information when analysis is complete
disp('   ')
disp('________________________________________________________________')
disp('   ')
disp('PARAMETERS FOR INTERESTING STRAINS (control is first)...')
disp('   ')
highlightsample = highlightsample(:);
if ~isempty(platesubset)
    if isempty(highlightsample)
        highlightinternalind = [];
    end
    disp(T(sort([internalind ; highlightinternalind]), :))
else
    disp(T(sort([controlstrainindex ; highlightsample]), :))
end
disp('________________________________________________________________')
disp('   ')
toc

disp('   ')

end
```
