## Supplementary File 2 for "Genomically recoded *Escherichia coli* with optimized functional phenotypes": iC321_DT_Analysis.m

```
%% iC321 data analysis
% Colin Hemez
% GitHub: colinfromtherandomforest
% 2021

clear; close all

%% Baseline subtraction on representative LB and M9 data
disp(' ')
disp('Baseline subtraction analysis')
F1Bfile = 'iC321_F1B_LB_F1_1';
F1Cfile = 'iC321_F1C_M9_F1_1';
LBtable = readtable([F1Bfile,'.xlsx']);
M9table = readtable([F1Cfile,'.xlsx']);
LBdata = table2array(LBtable(:,(2:end)));
M9data = table2array(M9table(:,(2:end)));

res = 0.001;

% For each LB curve, subtract baseline and calculate doubling time
LBdim = size(LBdata);
time = 10:10:(LBdim(1)*10);
time = time';
maxstart = max(LBdata(1,:));
maxrangeLB = 0.001:res:maxstart;
DT_LB_baseline = zeros(length(maxrangeLB), LBdim(2));
disp(' ')
for i = 1:LBdim(2)
    disp(['Analyzing LB curve ',num2str(i)])
    curvei = LBdata(:,i);
    startingOD = curvei(1);
    baserange = 0.001:res:startingOD;
    for j = 1:length(baserange)
        if baserange(j) > startingOD
            DT_LB_baseline(j,i) = NaN;
        else
            curvej = curvei - startingOD + baserange(j);
            DTi = lnDoublingTime(curvej, time, 7, 'plotoff');
            DT_LB_baseline(j,i) = DTi;
        end
    end
end
writematrix([maxrangeLB' DT_LB_baseline], 'OUTPUT_BaselineSubtraction_iC321_F1B_LB_F1_1', 'FileType', 'spreadsheet')

% For each M9 curve, subtract baseline and calculate doubling time
M9dim = size(M9data);
time = 10:10:(M9dim(1)*10);
time = time';
maxstart = max(M9data(1,:));
maxrangeM9 = 0.001:res:maxstart;
DT_M9_baseline = zeros(length(maxrangeM9), M9dim(2));
disp(' ')
for i = 1:M9dim(2)
    disp(['Analyzing M9 curve ',num2str(i)])
    curvei = M9data(:,i);
    startingOD = curvei(1);
    baserange = 0.001:res:startingOD;
    for j = 1:length(baserange)
        if baserange(j) > startingOD
            DT_M9_baseline(j,i) = NaN;
        else
            curvej = curvei - startingOD + baserange(j);
            DTi = lnDoublingTime(curvej, time, 7, 'plotoff');
            DT_M9_baseline(j,i) = DTi;
        end
    end
end
writematrix([maxrangeM9' DT_M9_baseline], 'OUTPUT_BaselineSubtraction_iC321_F1C_M9_F1_1', 'FileType', 'spreadsheet')

% Plot curves
figure
subplot(1,2,1)
plot(maxrangeLB, DT_LB_baseline, 'linewidth', 3)

subplot(1,2,2)
plot(maxrangeM9, DT_M9_baseline, 'linewidth', 3)

%% Clear workspace
clear;

%% Figure 1B/C

% Figure 1B: Standard strains in LB
F1Bfile = 'iC321_F1B_LB_F1_1';
ssOD = 0.030;
[~] = dataProcessing(F1Bfile, ssOD);

% Figure 1C: Standard strains in M9
F1Cfile = 'iC321_F1C_M9_F1_1';
ssOD = 0.020;
[~] = dataProcessing(F1Cfile, ssOD);

%% Figure 1D
F1Dfile = 'iC321_F1D_M9_F1_4';
ssOD = 0.020;
[~] = dataProcessing(F1Dfile, ssOD);

%% Supplementary Figure 2
FS2file = 'iC321_FS2_LB_F1_4';
ssOD = 0.030;
[~] = dataProcessing(FS2file, ssOD);

%% Figure 1F
F1Ffile = 'iC321_F1D_M9withsupplements_F1_3';
ssOD = 0.020;
[~] = dataProcessing(F1Ffile, ssOD);

%% Figure 2C
F2CFile = 'iC321_F2C_M9_F2_1';
ssOD = 0.020;
[~] = dataProcessing(F2CFile, ssOD);

%% Figure 2E
F2Efile1 = 'iC321_F2E_M9_F2_3';
F2Efile2 = 'iC321_F2E_M9_F2_4';
F2Efile3 = 'iC321_F2E_M9_F2_5';
ssOD = 0.020;
[~] = dataProcessing(F2Efile1, ssOD);
[~] = dataProcessing(F2Efile2, ssOD);
[~] = dataProcessing(F2Efile3, ssOD);

%% Supplementary figure 5
FS5file1 = 'iC321_FS5_LB_F2_3';
FS5file2 = 'iC321_FS5_LB_F2_4';
FS5file3 = 'iC321_FS5_LB_F2_5';
ssOD = 0.030;
[~] = dataProcessing(FS5file1, ssOD);
[~] = dataProcessing(FS5file2, ssOD);
[~] = dataProcessing(FS5file3, ssOD);
```
