## Supplementary File 2 for "Genomically recoded *Escherichia coli* with optimized functional phenotypes": lnDoublingTime.m

```
%% Calculation of doubling time by log2 transform
% Colin Hemez
% GitHub: colinfromtherandomforest
% 2021

function [DT,R,median] = lnDoublingTime(ODvals,timevals,points,varargin)

% Check that the number of points for linear fitting is odd
if mod(points, 2) ~= 1
    error('Subset of points for linear regression must be odd')
end

% Check that the number of values in OD and time arrays is equal
if length(ODvals) ~= length(timevals)
    error('ODVALS and TIMEVALS must have the same number of values')
end

plotter = 0;
datapts = length(ODvals);

if nargin == 4
    if strcmp(varargin{1}, 'ploton')
        disp('PLOT function active')
        plotter = 1;
        figure
    elseif strcmp(varargin{1}, 'plotoff')
        plotter = 0;
    else
        error('Optional fourth argument is invalid')
    end
end

% Transform the data into logspace, transform inf to NaN
ODlog = log2(ODvals);
ODlog(isinf(ODlog)) = NaN;
ODlog = real(ODlog);

% Perform linear regression to estimate doubling time
minRsq = 0.75; % R-squared value threshold
lowRsq = 0.90; % R-squared values below this threshold trigger a warning
padding = floor(points/2); % Number of points to consider on either side
maxSlp = 0; % Current highest slope
currVals = [0 0 0 0];

% currRsq: R-squared value of current highest slope

for i = (1+padding):(datapts-padding)
    Tmin = i - padding;
    Tmax = i + padding;
    currTvals = timevals(Tmin:Tmax);
    currODvals = ODlog(Tmin:Tmax);
    
    % Check if the region is above the linearity threshold
    currC = corrcoef(currTvals, currODvals);
    currR = currC(1,2);
    currRsq = currR^2;
    
    % If region is above linearity threshold, calculate the slope
    if currRsq >= minRsq
        linfit = polyfit(currTvals, currODvals, 1); % linfit: [slope, y-int]
        currSlp = linfit(1);
        
        % Update maximum slope; keep 
        if currSlp > maxSlp
            maxSlp = currSlp;
            Yint = linfit(2);
            currVals = [maxSlp Yint currRsq i];
        end
    end
    
end

maxVals = currVals;

% Do transformation on maximum values to get doubling time, centroid time
DT = 1 / maxVals(1);
R = maxVals(3);
if maxVals(4) > 0
    median = timevals(maxVals(4));
else
    median = 0;
end

% Trigger a warning if R-squared value is low but not terrible
if R < lowRsq
    warning(['Low R-squared value for sample. Rsq = ', num2str(R)])
end

% Trigger a warning if R-squared value is below threshold
if R < minRsq
    warning('R-squared value for sample is very low. Result unreliable.')
end

% Optional plotting function
if plotter == 1
    subplot(1,2,1)
    plot(timevals, ODvals, 'linewidth', 3)
    grid on
    xlabel('Time')
    ylabel('OD_{600}')
    title('Raw Data')
    
    subplot(1,2,2)
    hold on
    
    % Plot log-transformed data
    plot(timevals, ODlog, 'ok', 'markerfacecolor','k')
    
    % Plot the region of linear fit used to calculate doubling time
    fitTimes = timevals( (maxVals(4)-padding) : (maxVals(4)+padding) );
    fitLnOD = (fitTimes.*maxVals(1)) + maxVals(2);
    plot(fitTimes, fitLnOD, '-r', 'linewidth', 5)
    
    grid on
    xlabel('Time')
    ylabel('log_2 (OD_{600})')
    title('Log-Transformed Data')
end

end
```
