## Supplementary File 2 for "Genomically recoded *Escherichia coli* with optimized functional phenotypes": README_iC321_SuppFile2.txt

This folder contains the MATLAB scripts used to process growth curve data reported in Hemez et al., "Genomically recoded Escherichia coli with optimized functional phenotypes."

iC321_DT_Analysis: Script that analyzes all growth curves for doubling time and maximum OD, and that performs baseline subtraction analysis on representative LB and M9 growth curve data. This file calls growthCurveAnalysis to determine doubling times and maximum ODs, and can be used to process the input data files given in Supplementary file 1.

dataProcessing: Runs growthCurveAnalysis on an Excel data file formatted with timepoints in the first column and sample names in the first row.

growthCurveAnalysis: Function that calculates doubling time maximum OD for an input file containing raw OD values. growthCurveAnalysis calls lnDoublingTime to calculate doubling times on individual growth curves.

lnDoublingTime: Calculates doubling time for an individual growth curve. This function transforms a growth curve time series into logarithmic space and determines the steepest linear portion of the log-transformed curve over a specified number of timepoints (default is 7). The slop of this region is used to calculate doubling time.

All data were processed using MATLAB release R2019a.
